# Brain-predicted age associates with psychopathology dimensions in youth

**DOI:** 10.1101/2020.06.13.149658

**Authors:** Vanessa L. Cropley, Ye Tian, Kavisha Fernando, L. Sina Mansour, Christos Pantelis, Luca Cocchi, Andrew Zalesky

## Abstract

**Background:** This study aims to investigate whether dimensional constructs of psychopathology relate to advanced, attenuated or normal patterns of brain development, and to determine whether these constructs share common neurodevelopmental profiles.

**Methods:** Psychiatric symptom ratings from 9312 youths (8-21 years) were parsed into 7 independent dimensions of clinical psychopathology representing conduct, anxiety, obsessive-compulsive, attention, depression, bipolar, and psychosis symptoms. Using a subset of this cohort with structural MRI (*n*=1313), a normative model of brain morphology was established and the model was then applied to predict the age of youth with clinical symptoms. We investigated whether the deviation of brain-predicted age from true chronological age, called the brain age gap, explained individual variation in each psychopathology dimension.

**Results:** Individual variation in the brain age gap significantly associated with clinical dimensions representing psychosis (*t*=3.16, *p*=0.0016), obsessive-compulsive symptoms (*t*=2.5, *p*=0.01), and general psychopathology (*t*=4.08, *p*<0.0001). Greater symptom severity along these dimensions was associated with brain morphology that appeared older than expected for typically developing youth of the same age. Psychopathology dimensions clustered into two modules based on shared brain loci where putative accelerated neurodevelopment was most prominent. Patterns of morphological development were accelerated in frontal cortices for depression, psychosis and conduct symptoms (Module I), whereas acceleration was most evident in subcortex and insula for the remaining dimensions (Module II).

**Conclusions:** Our findings suggest that advanced brain development, particularly in frontal cortex and subcortical nuclei, underpins clinical psychosis and obsessive-compulsive symptoms in youth. Psychopathology dimensions share common neural substrates, despite representing clinically independent symptom profiles.

## Introduction

Childhood and adolescence is marked by profound changes in cerebral gray and white matter structure. From middle childhood to early adulthood, gray matter volume decreases, whereas white matter increases (1, 2), though precise developmental trajectories vary across regions, structural measures and in their timing (3–6). Many mental illnesses first manifest during this period of rapid neurodevelopment. Often considered as neurodevelopmental disorders, mental illness is also known to affect brain structure and function (7).

Both *delayed* and *advanced* patterns of gray matter development associate with psychiatric symptoms in youth. Longitudinal studies report *delayed* or *attenuated* cortical maturation in children with attention deficit hyperactivity disorder (ADHD) (8), and in youth with internalizing (9, 10), and externalizing (10) symptoms. Conversely, *increased* cortical thinning or volume loss associates with depression in adolescence (11), and over the transition to and onset of psychosis (12–15). Aberrant development of subcortical volumes (11, 16, 17) and white matter (16, 18), also relate to psychopathology, though the specific changes vary across studies. These findings suggest that *deviation* from normative patterns of brain development is important in the emergence of psychopathology in youth.

The extent to which an individual deviates from a healthy neurodevelopmental trajectory can be quantified with normative models that aim to predict an individual’s age based on their brain scans. The *brain age gap* is defined by subtracting chronological age from the brain-predicted age inferred from a normative model (19–21). A person with a positive brain age gap has a brain that appears to be older than the person’s chronological age, whereas a negative brain age gap implies that the person’s brain appears younger than people of the same age.

Brain age gap is increasingly examined in individuals with psychiatric disorders, principally in adult populations (22). Advanced brain age scores have been reported in major depressive disorder (23), schizophrenia (22–26), individuals at clinical high risk for developing psychosis (23, 27) and bipolar disorder in some (22) but not all (24, 25) studies. While these studies suggest that the brain appears older than its chronological age in most adult patient cohorts, the neurodevelopmental period at which deviations between chronological and brain age emerge remains unclear. Here, we investigated the brain age gap across multiple dimensions of psychopathology in youth, aiming to understand whether specific clinical symptoms were marked by deviations in brain development during a crucial developmental period. We also aimed to determine whether the brain loci with the most prominent deviations between chronological and brain-predicted age were shared between independent psychopathology dimensions.

To address these aims, we established a normative model of brain age based on brain morphology derived from structural MRI acquired in a large sample of youths (*n*=1313) aged 8 to 21 years comprising the Philadelphia Neurodevelopmental Cohort (PNC). We investigated whether the deviation between brain-predicted and chronological age associated with 7 dimensions of psychopathology measuring the clinical severity of conduct, anxiety, obsessive-compulsive, attention, depression, bipolar and psychosis symptoms, as well as overall psychopathology. We then tested whether these independent dimensions could be stratified based on shared patterns of putative accelerated or delayed neurodevelopment. We expected that (sub)clinical psychiatric phenomena would be associated with subtle deviations in normative brain development, as indexed by the brain age gap, and hypothesized that the psychopathology dimensions would form distinct clusters based on common patterns of altered neurodevelopment. This study provides new insights into early brain changes that may lead to psychiatric symptoms and identifies a common neural basis for several clinically distinct dimensions of psychopathology.

## Methods and Materials

### Participants

Recruitment procedures, sample characteristics and assessment protocols for the PNC are described elsewhere (28) and in Supplement 1.1. The Institutional Review Boards of the University of Pennsylvania and the Children’s Hospital of Philadelphia approved all study procedures. All participants provided written informed consent as well as parental/legal guardian permission for participants under 18. Of 1598 participants for whom a structural MRI brain scan was acquired, 285 participants were excluded due to presence of severe medical conditions (*n*=74), image quality control (*n*=209; see Supplement) or missing clinical data (*n*=2), leaving a total of 1313 youths (49.8% male) aged 8-21 (mean 14.5, SD 3.43 years). Within this final sample, 402 individuals were identified as typically developing (TD) control youths, operationalized here as having no medical conditions that could affect the central nervous system, no history of psychiatric hospitalization, not currently taking psychotropic medication and not meeting criteria for psychosis spectrum symptoms (29). The remaining 911 youth were not confirmed to meet the above constraints and were thus classified as non-TD (see Supplement 1.1). **Supplementary Table 1** presents demographic characteristics of the study sample.

### Assessment of psychopathology

Assessment of lifetime psychopathology was performed using a structured screening interview (GOASSESS) (28). Full details are described in Supplement 1.2. Collateral informants were used for participants 8-10 years of age. Independent component analysis (ICA) was used to factorize the 129 symptom scores (**Supplementary Table 2**) comprising the GOASSESS into dimensional measures of psychopathology, as has been previously validated in the PNC (30). ICA was performed on the full PNC sample, including participants without MRI data (*n*=9312) to obtain stable independent components using the largest possible sample size (see Supplementary 1.3). This yielded 7 independent components representing continuous dimensions of clinical symptoms related to *Conduct*, *Anxiety*, *Obsessive-Compulsive*, *Attention*, *Depression*, *Bipolar*, and *Psychosis*. Each individual was thus represented by 7 scores indexing severity for each of these dimensions, from symptom absence (lowest score) to clinical psychopathology (highest score). A measure of *General Psychopathology* was defined by averaging the 7 scores for each individual, yielding 8 psychopathology measures in total. As this study adopted a dimensional approach to evaluating psychopathology, no threshold psychiatric diagnoses were established for any participant. The sample of 1313 youths (consisting of TD and non-TD individuals) with both structural MRI and ICA scores were subsequently analyzed.

### Image acquisition, processing and quality control

A 3 Tesla Siemens MRI scanner was used to acquire a T1-weighted (MPRAGE) structural MRI scan with the following parameters: TR: 1810 ms; TE: 3.51 ms; FOV: 180 × 240 mm; matrix: 256 × 192; 160 slices; TI: 1100 ms; flip angle: 9°; effective voxel resolution: 0.9 × 0.9 × 1 mm.

Image processing was conducted using Freesurfer v6.0 (31) to estimate cortical surfaces and subcortical segments for each participant. Seven subcortical volumes (nucleus accumbens, amygdala, caudate, hippocampus, pallidum, putamen, and thalamus), 1 lateral ventricle volume, ICV and 34 cortical regions for each of cortical thickness, surface area and volume (111 measures in total) were extracted as features for brain age prediction. Prior to feature extraction, automated quality assessment of T1 scans was performed using Freesurfer’s Euler number and the Mahalanobis distance of cortical thickness measurements (32, 33). Full details are found in Supplement 1.4-1.5.

### Prediction of age and calculation of brain age gap

Using 10-fold cross-validation on the 402 TD youth (49% male), a linear support vector regression (SVR) model was trained to predict individual age based on 111 regional measurements of gray matter volume, thickness and area. All measures were mean and variance standardized. The predicted age for each TD individual was determined using the model that was trained for the fold in which the individual comprised the held-out set. Model performance was quantified by calculating the mean absolute error (MAE) and the Pearson correlation coefficient between predicted age and chronological age. In supplementary analyses, the model was trained on the full sample using 10-fold cross validation, rather than only the TD group, to assess the impact of sample stratification. Individual age was also predicted using a multi-modal feature space comprising the 111 grey matter measures and 192 regional diffusion measures of white matter integrity (fractional anisotropy, mean, radial and axial diffusivity; see Supplement 1.6 for details). The brain age gap was calculated for each individual by subtracting chronological age from the brain-predicted age. Based on recent recommendations (34, 35), we regressed out the effect of age on brain age gap in order to correct for the ‘regression to the mean’ bias and used the resulting residuals in further analyses. The overall methodology is schematized in **Supplementary Figure 1**.

### Association of brain age gap with psychopathology

A general linear model was formulated to test whether brain age gap associated with each of the dimensions of psychopathology in the non-TD youth (*n*=911). The brain age gap of the *i*th participant was modeled as,

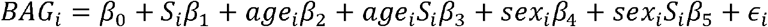

where *BAG*_*i*_ and *S*_*i*_ denote the residualized brain age gap (34) and psychopathology score of the *i*th participant, respectively, *β* denotes the fitted regression coefficients and *ϵ*_*i*_ is the error term. The model was fitted independently to each of the 8 psychopathology scores. The explanatory variable characterizing sex (*β*_4_), and the age-by-psychopathology (*β*_3_) and sex-by-psychopathology (*β*_5_) interactions were not significant and hence omitted from the final model. The false discovery rate (FDR) was controlled at 5% across the 8 independent tests.

### Regional localization of association between brain age gap and psychopathology

To identify the cerebral loci that were most informative to the relation between brain age gap and psychopathology, the SVR model was successively re-trained with features pertaining to specific brain lobes (i.e. frontal, parietal, temporal, occipital, cingulate, insula, subcortical) or regions (i.e. lateral ventricle, ICV) omitted. Formation of cortical lobes followed the standard Desikan-Killiany Freesurfer convention (https://surfer.nmr.mgh.harvard.edu/fswiki/CorticalParcellation).

Specifically, we computed the strength of the association between individual variation in brain age gap and psychopathology scores after each of the 9 lobes/regions was omitted from the feature set used to train the SVR. This yielded a 9 × 8 matrix (lobes/regions × psychopathology dimensions) of *t*-statistic values quantifying the strength of the associations between psychopathology and brain age gap computed with information from a specific lobe/region missing. The importance of a lobe/region was quantified by the decrease in the strength of the association following omission of the lobe/region from the SVR.

### Clustering of psychopathology dimensions

A clustering analysis was performed to determine whether a shared pattern of brain morphology existed across the psychopathology dimensions. The Pearson correlation coefficient was computed between all pairs of columns of the 9 × 8 matrix of *t*-statistics described above, yielding an 8 × 8 correlation matrix (dimensions × dimensions). Correlation coefficients were highest between psychopathology dimensions for which the same brain lobes/regions were most informative to the association between psychopathology and brain age gap. The rows/columns of the correlation matrix were reordered to reveal two unequivocal clusters among the psychopathology dimensions. Ward’s clustering method was applied to the 9 × 8 matrix of *t*-statistics to confirm the clusters.

### Associations between subcortical and cortical brain structures with psychopathology

To align with previous studies using classical group-level inference, supplementary analyses tested the extent to which each psychopathology dimension accounted for individual variation in cortical surface measures, and subcortical volumes, controlling for age and sex. The spin test (36) was used to test whether the areas of the cortex where the relationship between cortical structure and psychopathology was strongest/weakest were spatially consistent across the dimensions (Supplement 1.8).

## Results

In a large sample of youth, we identified 7 continuous dimensional constructs of clinical psychopathology (**Figure 1A**). For each dimension, individuals were scored on a spectrum ranging from absence of symptoms (lowest score), to subclinical and possibly threshold symptoms (highest score). The psychopathology dimensions were found to cut across classical diagnostic categories (**Figure 1B**), although clinical diagnoses were not established for any participant. Our dimensional constructs of psychopathology are consistent with Alneas and colleagues (30), although others (37, 38) provide a complementary set of dimensions. Supplement 2.1 describes demographic and clinical characteristics.

**Figure 1.**
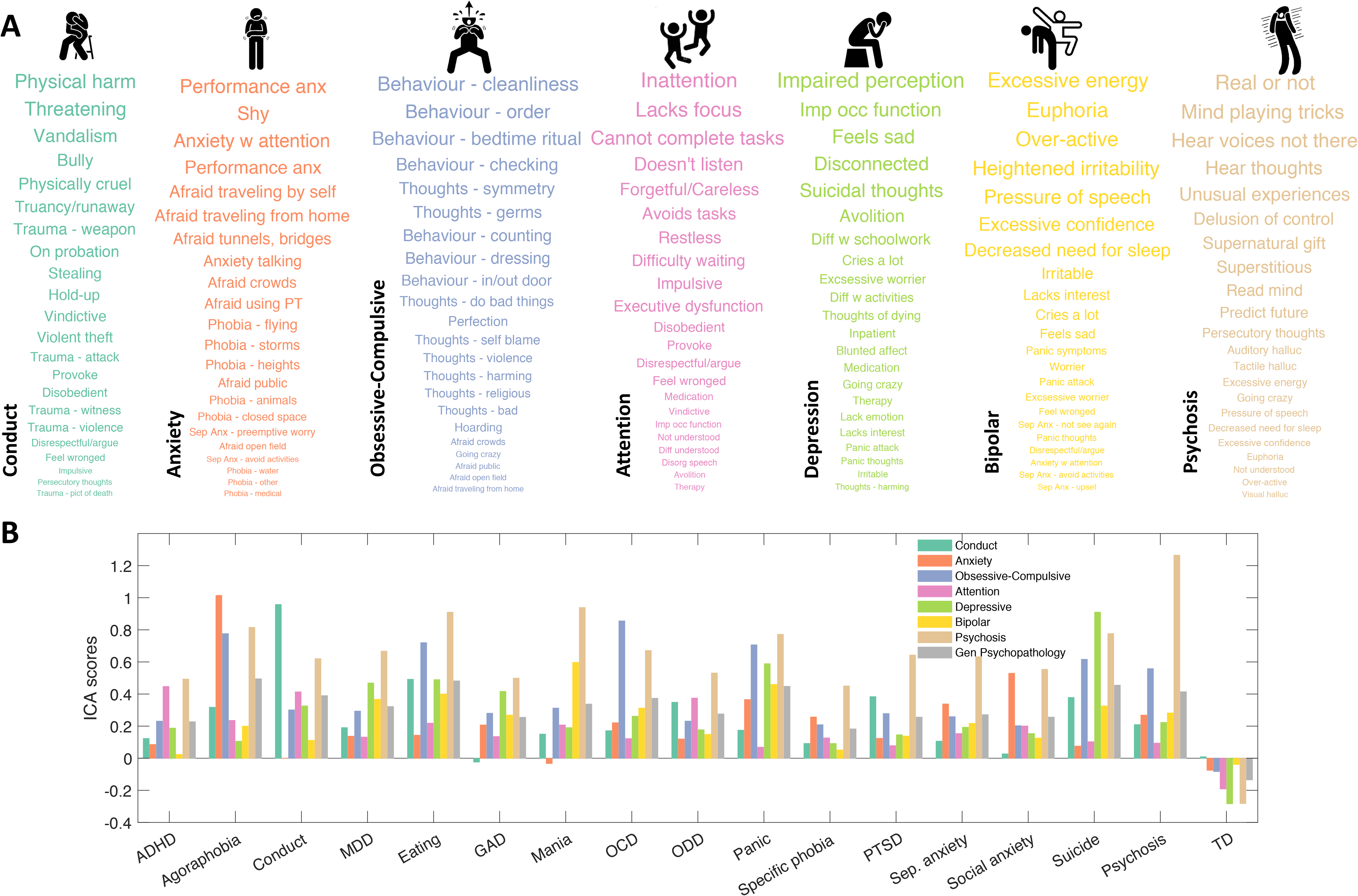
Independent dimensions of clinical psychopathology in a large youth cohort. (**A**). Independent component analysis (ICA) was used to identify continuous dimensions of psychopathology in a large sample of youth (n=9312). The 129 items comprising the GOASSES interview were parsed into 7 independent dimensions of psychopathology representing: i) Conduct, ii) Anxiety, iii) Obsessive-Compulsive, iv) Attention, v) Depression, vi) Bipolar, and vii) Psychosis. The GOASSES items that loaded most strongly onto each dimension are shown. Font size is scaled according to the absolute value of loading strength. Thus, the GOASSES items most representative of a dimension are shown in the largest font sizes. Icons are taken from https://www.flaticon.com/. (**B**). Bar plots show the average score for each of the 7 psychopathology dimensions, stratified by conventional diagnostic categories. Established diagnostic categories are shown on the horizontal axis. Each bar represents the average psychopathology score across youth comprising the diagnostic category for the subsample with neuroimaging data (n=1313). Specifically, non-TD individuals who endorsed any of the symptom items pertaining to each of the 16 clinical screening categories were identified, yielding 16 groups of individuals with lifetime endorsement of these clinical categories. Note that the two screening categories of prodromal symptoms and general probes were not included. Some individuals were assigned to multiple clinical groups, whereas TD individuals formed a single group. Each of the 7 component scores and the score of general psychopathology were then averaged across individuals within each diagnostic category. ADHD, attention-deficit hyperactivity disorder; MDD, major depressive disorder; GAD, generalized anxiety disorder; OCD, obsessive-compulsive disorder; ODD, oppositional defiance disorder; PTSD, post-traumatic stress disorder; TD, typically developing.

### Predicting age based on brain morphology

For the typically developing (TD) group that was used to train the normative model in the first instance, structural brain measures predicted chronological age with a mean absolute error (MAE) of 1.49 years (average over 100 cross-validation partitions) and a correlation between chronological and predicted age of *r*=0.82, *p*<0.0001. Applying the trained model to the non-TD individuals yielded age predictions of comparable accuracy (MAE: 1.57 years, r=0.72, p<0.0001). The beta weights for each individual gray matter feature in the prediction of age are visualized in **Figure 2**. Gray matter features showed both positive and negative correlations with predicted brain age, which varied regionally and with gray matter metrics. Incorporating white matter diffusion measures (fractional anisotropy, mean, radial and axial diffusivity) into the normative model did not improve brain age prediction (Supplement 2.2). These measures were thus given no further consideration here.

**Figure 2.**
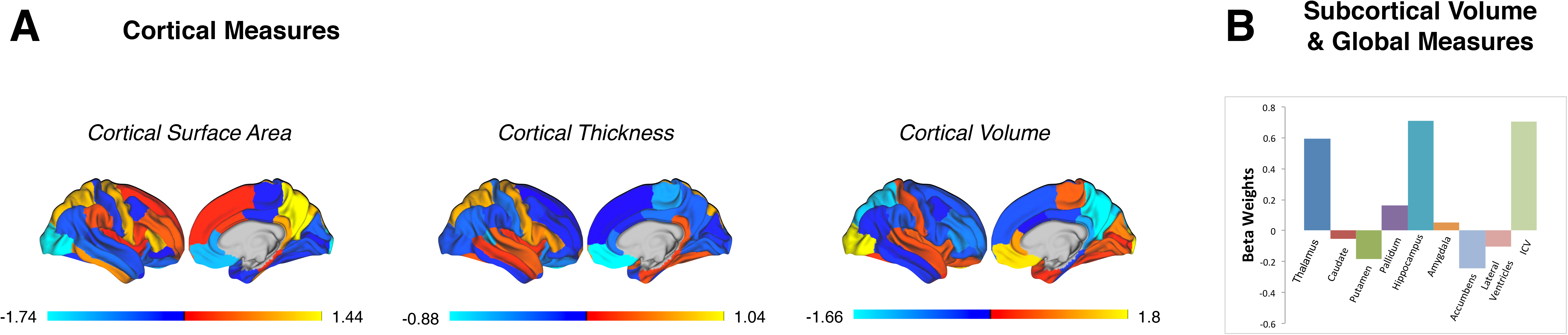
Relative importance of gray matter features in predicting age. (**A**) Visualized are the beta weights for each individual cortical region for surface area, thickness and volume in the prediction of age. Cortical regions are based on the Desikan-Killiany parcellation atlas. Warmer colours correspond to increased cortical morphology with increasing age, whereas cooler colours correspond to decreased morphology with increasing age. Only one hemisphere is shown due to the averaging of left and right hemispheres for each region. (**B**) Bar plot shows the beta weights for each individual subcortical volume in the prediction of age. Positive beta weights correspond to increases in volume with increasing age, whereas negative beta weights correspond to decreases in volume with increasing age.

### Association between brain age gap and psychopathology dimensions

Individual variation in brain age gap was significantly associated with dimensions representing obsessive-compulsive symptoms (OCS; *t*=2.5, *p*=0.01) and psychosis (*t*=3.16, *p*=0.0016) as well as general psychopathology (*t*=4.08, *p*<0.0001) (**Figure 3**). Individuals with brain morphology that appeared *older* than predicted by the normative model were associated with *greater* symptom severity along these dimensions. Brain age gap was not associated with any of the other psychopathology dimensions (*p*>0.05).

**Figure 3.**
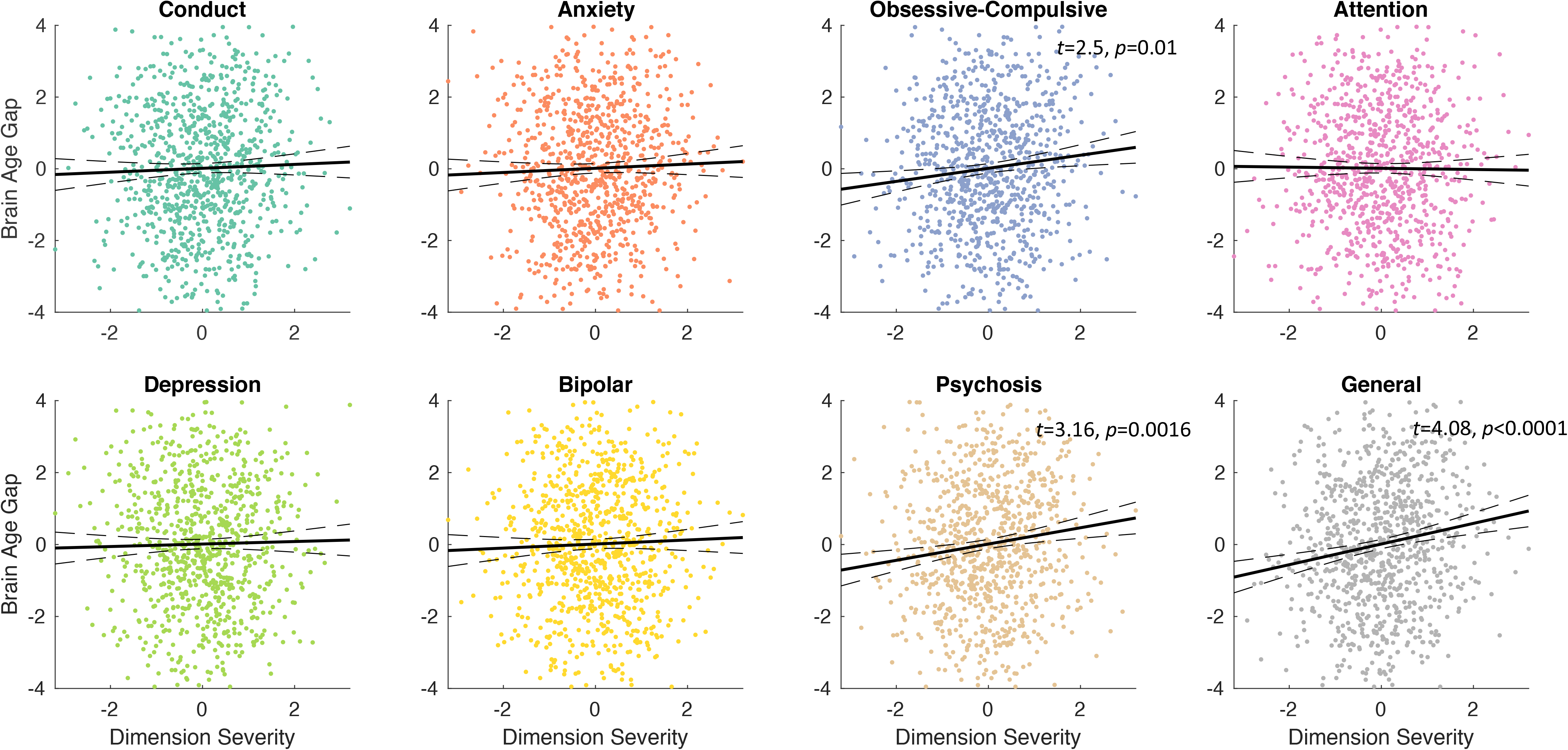
Associations between brain age gap and individual variation in dimensions of psychopathology. Individual variation in the difference between chronological age and brain-predicted age (brain age gap) was significantly associated with subclinical dimensions representing psychosis (*t*=3.16, *p*=0.0016), OCS (*t*=2.5, *p*=0.01), and general psychopathology (*t*=4.08, *p*<0.0001). The higher an individual scored on a symptom dimension, the older their brain morphology appeared. Each data point represents an individual (n=1313). Lines of best fit (solid) and 95% confidence intervals (dashed) are shown. A scatter plot is shown for each of the 7 psychopathology dimensions tested as well as the measure of general psychopathology, whereas test statistics and p-values are only shown for the dimensions significantly associated with brain age gap, controlling the false discovery rate (FDR) at 5%.

To confirm these findings, we tested whether any of the 129 GOASSES items associated with brain age gap. Thirty-seven of the items were significantly related to brain age gap (FDR<5%), with the majority of significant associations relating to psychosis and OCS (**Supplementary Table 3**). We also computed average brain age gap for each dimension by computing a weighted mean over all non-TD individuals, where each individual’s brain age gap was weighted by their psychopathology score. Average brain age gap was significantly greater than zero for general psychopathology (76 days older than predicted by the normative model), OCS (66 days older) and psychosis symptoms (52 days older), indicating an older than expected brain age for each of these dimensions (**Supplementary Table 4**).

To further validate the above relationships, we substituted the continuous psychopathology dimensions with a categorical operationalization of psychopathology. Individuals with a lifetime endorsement of symptoms pertaining to the psychosis and obsessive-compulsive screening categories in the GOASSESS were grouped into a subclinical psychosis (*n*=328) and subclinical OCS (*n*=391) group, respectively (**Supplementary Table 5**). Brain age gap was significantly higher in the psychosis (*t*=2.35, *p*=0.02) and OCS (*t*=2.35, *p*=0.021) groups compared to the TD group. The interaction between age and group was also significant (psychosis: *p*<0.001; OCS: *p*=0.002), indicating that the brain age gap was largest for the youngest individuals in the cohort who expressed psychosis and OCS symptoms (**Figure 4**). Brain age gap did not differ between the TD and non-TD groups (**Supplementary Figure 2**).

**Figure 4.**
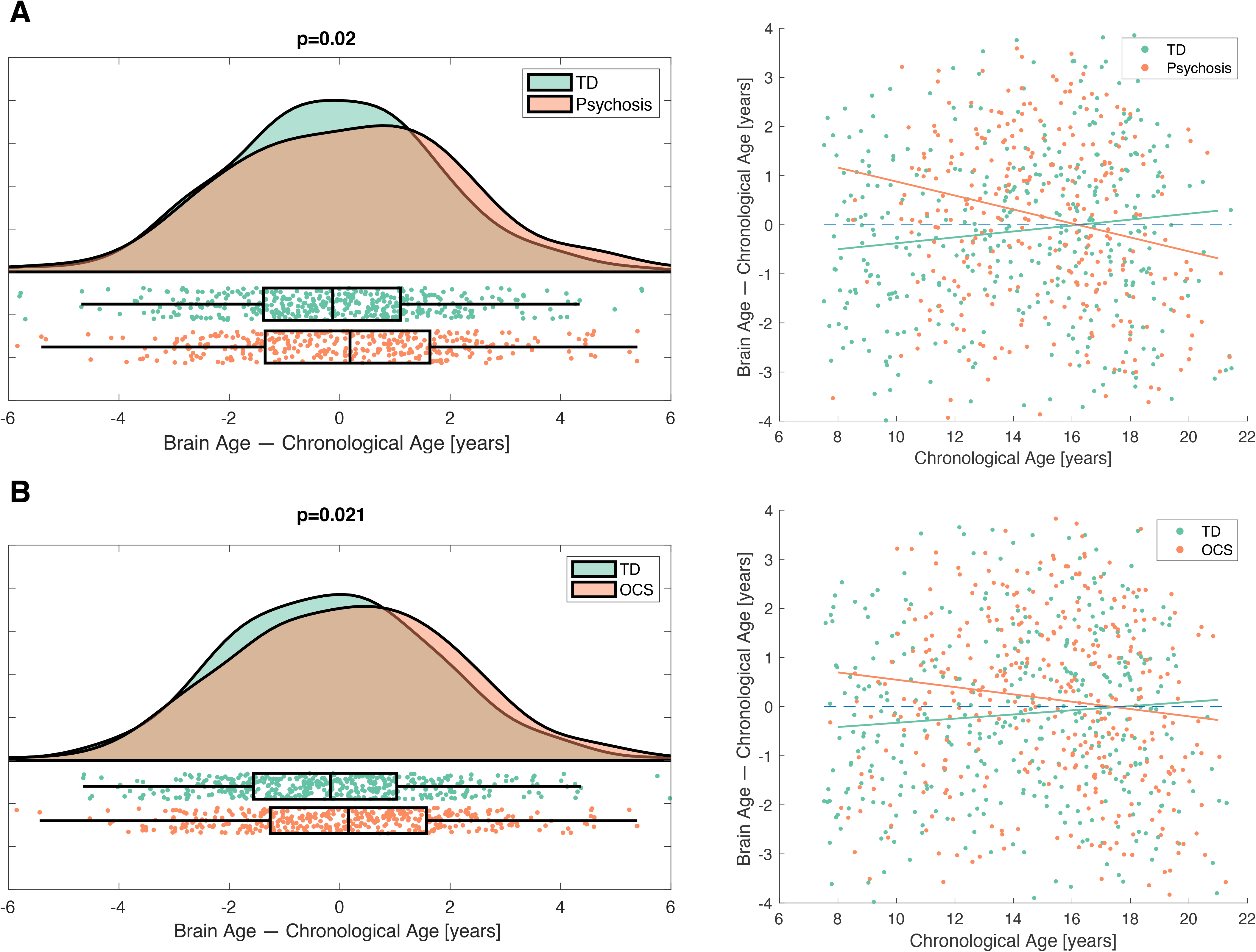
Advanced brain aging in individuals with endorsement of subclinical psychosis and obsessive-compulsive symptoms. The difference between chronological age and brain-predicted age (brain age gap) was significantly increased in youth endorsing psychosis (**A**) and OCS (**B**) relative to TD youth. Categorical groupings were formed on the basis of endorsement of any item within the psychosis and obsessive-compulsive GOASSES screening categories. Box plots (left) show the distribution of brain age gap for the TD (green) and the two non-TD groups (red). An age-by-group interaction was significant for both categories (psychosis: *p*<0.001; OCS: *p*=0.002), represented by greater brain age gap in younger individuals endorsing psychosis and OCS. Scatter plots (right) show the age-by-group interaction with lines of best fit shown for the TD (green) and two non-TD groups (red). OCS, obsessive-compulsive symptoms; TD, typically developing.

Supplementary analyses were undertaken to establish the replicability and robustness of the associations between brain age gap and psychopathology. First, all the associations remained significant and were replicated in the entire cohort (i.e. TD and non-TD combined). This established that the associations were not contingent on the operationalization of TD. Second, to assess the impact of sample dichotomization, the SVR model was re-trained based on the entire cohort using 10-fold cross-validation, rather than only on the non-TD individuals. All associations were once again replicated. This established that the associations were robust to both categorical (i.e. TD/non-TD) and dimensional characterizations of the cohort. Finally, all the associations remained significant after controlling for potential confounds, including medical rating, psychotropic medication use, IQ, and traumatic-stress exposure. Traumatic-stress exposure associated with psychopathology and showed an interaction with age on brain age gap. Details are provided in Supplementary 1.7 and 2.3.

### Brain regions mediating the association between brain age gap and psychopathology

To determine which brain lobes/regions were most informative to the above associations, we systematically omitted each specific lobe/region from the normative model and quantified the extent to which this weakened the association between brain age gap and psychopathology. The greatest reduction in variance explained for psychosis was observed with frontal lobe exclusion, whereas the greatest reduction in variance explained for OCS and general psychopathology was found when excluding subcortical volumes, and the insula, respectively (**Figure 5A & B**).

**Figure 5.**
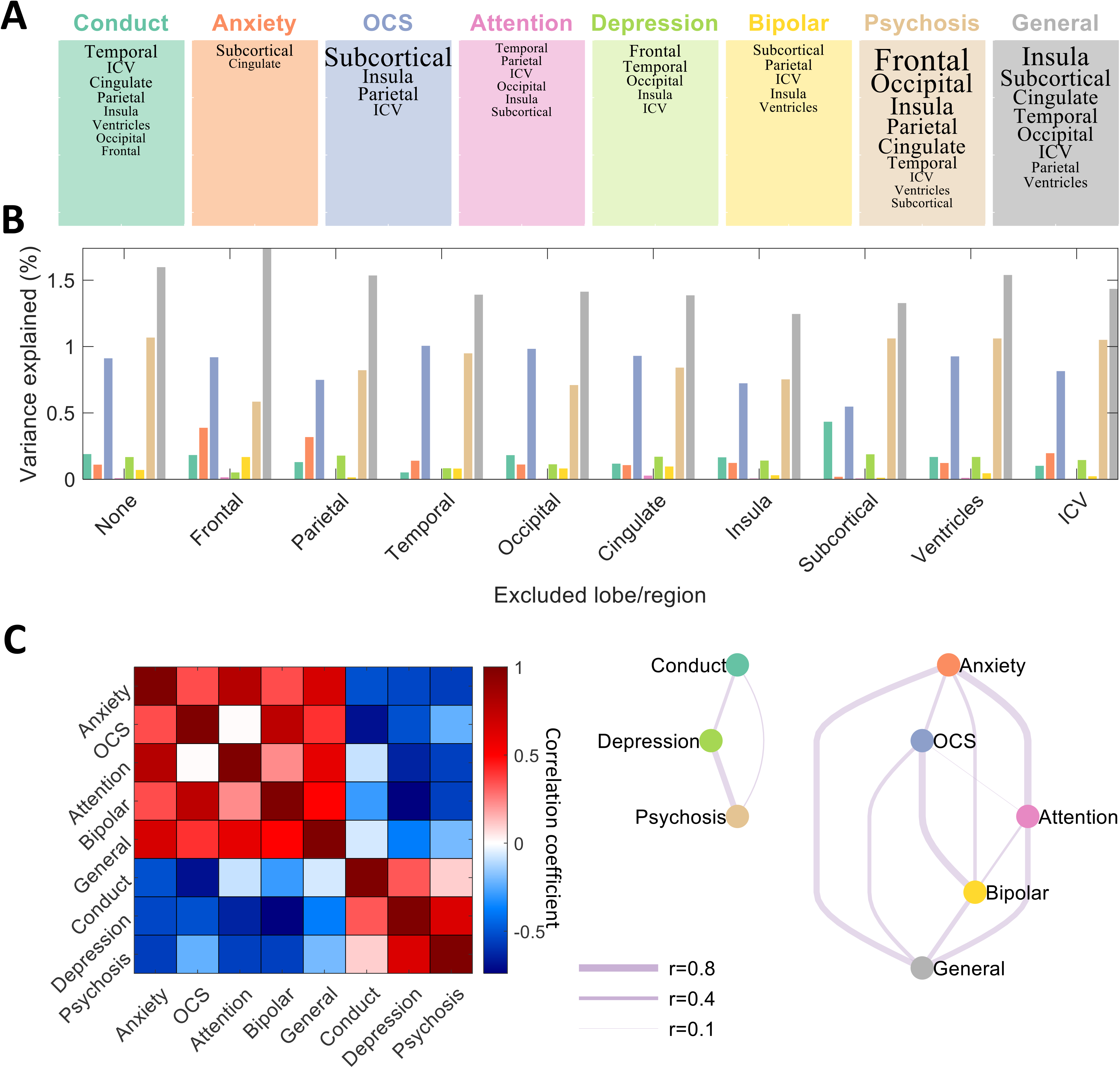
Clustering of psychopathology dimensions based on common neural substrates. The omission of lobes/regions weakened the association between brain age gap and psychopathology differently across the psychopathology dimensions. (**A**) The features that reduced the strength of the association between psychopathology and brain age gap are shown for each of the 7 dimensions as well as general psychopathology. Text in larger font indicates a greater relative reduction in effect size. (**B**) Reduction in variance explained by each of the 9 feature omissions is depicted for each of the psychopathology dimensions. (**C**) The correlation matrix (left) and corresponding network representation (right) show stratification of the psychopathology dimensions into two distinct clusters (modules) based on common neural substrates. Altered morphological development was circumscribed to the same brain regions/lobes for psychopathology dimensions comprising the same cluster. One cluster (Module I) comprises psychosis, depression and conduct dimensions, whereas general psychopathology together with the OCS, anxiety, bipolar and attention dimensions form the second cluster (Module II). Module I was characterized by accelerated morphological development in frontal cortices, whereas this acceleration was most evident in the subcortex and insula for Module II. OCS, obsessive-compulsive symptoms.

### Independent psychopathology dimensions share common neural substrates

Follow-up analyses were performed to determine whether the psychopathology dimensions formed distinct clusters based on shared patterns of brain morphology that were most informative to explaining individual variation in psychopathology. Two distinct clusters (modules) of dimensions were evident (**Figure 5C**). One cluster comprised psychosis, depression and conduct dimensions (Module I), whereas general psychopathology together with the OCS, anxiety, bipolar and attention dimensions formed a distinct cluster (Module II). Each cluster defined a distinct topographical pattern of accelerated cortical development. In particular, morphological development was accelerated in the cortex (and particularly the frontal cortex) for depression, psychosis and conduct symptoms (Module I), whereas acceleration was most evident in the subcortex and insula for the remaining dimensions (Module II). Despite not all of the psychopathology dimensions showing an association with brain age gap, these results reveal that the dimensions formed two distinct clusters based on shared regional patterns of accelerated development.

### Characterizing the relationship between dimensions of psychopathology and brain structure

Finally, we aimed to determine to what extent each psychopathology dimension associated with individual variation in brain morphology when controlling for age and sex. We found that increased severity of psychopathology related to lower subcortical and cortical structure within specific regions. These regions were common to the anxiety, conduct and psychosis dimensions (Supplement 2.4).

## Discussion

Childhood and adolescence is a critical period of brain maturation (39) and a risk period for the emergence of the first symptoms of mental illness (7). Using a large community sample of youth, we mapped dimensional constructs of clinical psychopathology and investigated whether the dimensions related to a novel index of neurodevelopment called the brain age gap (19–21). We found that the brain age gap increased with increasing severity of symptoms along the dimensions representing psychosis and OCS, as well as general psychopathology. The brain appeared older than its true chronological age in youth scoring highly on these dimensions, suggesting *accelerated* brain maturation. Moreover, we found that the psychopathology dimensions clustered into two distinct modules based on common brain loci where accelerated maturation was most prominent. Hence, while the dimensions were operationalized in such a way to ensure their independence with respect to symptomatology, several dimensions shared similar neurodevelopmental profiles, suggesting a common neuropathological basis.

Given the specificity of our findings to the psychosis and OCS dimensions, it is unlikely that greater gap scores were solely driven by severity of general psychopathology. Our findings were also robust to the choice of training sample, as well as select confounds, including psychotropic medication. Similar to Gur and colleagues (40), we found that greater traumatic-stress exposure was associated with greater severity of most dimensions, and, in younger individuals, subtle increases in brain age gap (Supplement). Importantly, the associations of brain age gap with OCS, psychosis and general psychopathology persisted when controlling for trauma and its interaction with age. This suggests that our measure of brain age gap is capturing unique variance in these dimensions, which is not explained by trauma alone. Examination of other forms of early life adversity, such as deprivation, is warranted.

Several processes can potentially explain why accelerated brain maturation was related specifically to OCS and psychosis dimensions, but not other dimensions. There is high comorbidity between OCD and schizophrenia, with evidence that OCD precedes schizophrenia onset (41). Recent work using the PNC found that subthreshold OCS was associated with higher rates of severe psychiatric conditions, including psychosis (42). In this study, endorsement of OCS representing “bad thoughts” had the greatest effect on major comorbid psychopathology. This finding highlights that particular OCS patterns may be a harbinger for other psychopathology, including psychosis. Notably, we demonstrated that OCS items reflecting negative intrusive thoughts were most strongly associated with greater brain age gap (see Supplement). Negative thought patterns might therefore be particularly associated with accelerated brain structural maturation and shared across OCS and psychosis dimensions.

The precise regions that showed accelerated maturation varied between the psychopathology dimensions. Nevertheless, we identified two characteristic structural patterns that separated the conduct, depression and psychosis dimensions, from the remaining dimensions. Regionally, these distinct patterns could be characterized by cortical (Module I) and subcortical (Module II) pathology. In particular, the accelerated pattern of structural development characterizing psychosis was primarily mediated by the frontal lobe, whereas for OCS and general psychopathology this pattern was mediated by the subcortex and insula. These brain regions have been linked to core pathological processes in psychosis (43) and OCD (44), respectively. Therefore, although variation in psychosis and OCS were both linked to increased brain age gap, the regions mediating this pattern were distinct. The mechanisms underlying brain age gap (and their spatial differences) are unknown and likely diverse, relating to lifestyle, environmental, substance use, biological and genetic ^((i.e. 22))^ factors, and may be shared or distinct across psychosis and OCS dimensions. In particular, excessive or premature synaptic pruning may be plausibly involved given its neurodevelopmental role (45) and implication in schizophrenia (46). Future studies are required to elucidate these mechanisms.

The current study accords with findings of greater brain age (22, 24–27), and cortical thinning/volume loss (12–15, 47), across the psychosis spectrum. Our findings also broadly accord with the recent work of Kaufmann et al (22) who adopted a transdiagnostic case-control approach. These authors found advanced brain aging in neurological disorders and schizophrenia, and to a lesser extent, in bipolar and psychosis-spectrum disorder. Our study extends this previous work by demonstrating subtle deviations in neurodevelopment with a dimensional construct of psychopathology that may have captured subclinical symptoms. Utilising a categorical approach we also revealed a significant group by age interaction indicating higher brain age gap in younger individuals endorsing psychosis or OCS. This finding is consistent with reports in clinical high-risk for psychosis individuals (27), and suggests that youth with an earlier onset of these symptoms may be particularly vulnerable to altered brain development.

Several limitations require consideration. Firstly, our brain age prediction model was based solely on gray matter features derived from a regional atlas. It is possible that multi-modal neuroimaging, and/or increased spatial scale, may increase the accuracy of brain age prediction. Nevertheless, we found that the addition of diffusion features did not improve age estimates (see Supplement). Secondly, while highly significant, the associations between brain age gap and psychopathology were modest in terms of effect size. Considering nonlinear relations could have potentially increased the variance explained and effect sizes, but introduces a risk of model overfitting. Finally, our study was based on an age-diverse cross-sectional design. While it may be tempting to infer that a positive brain age gap corresponds to acceleration in brain development or aging, whereas a negative age gap implies delayed or arrested development, such conclusions are not necessarily warranted in the absence of longitudinal measurements.

## Conclusions

In line with the Research Domain Criteria (RDoC) initiative (48) we adopted a dimensional approach to probe brain-psychopathology relationships and employed novel normative modelling to characterize individual heterogeneity in brain development. Our use of a large, community sample of youth allowed us to identify relationships between brain structure and a spectrum of psychopathology ranging from absent, to subclinical and threshold severity. Our findings suggest that accelerated development of brain structural morphology, particularly in frontal cortex and subcortical nuclei, may underpin psychosis and obsessive-compulsive symptoms in youth, particularly in individuals with an earlier onset of psychopathology. We further found that common patterns of altered brain development underpinned psychopathology constructs that are independent from a clinical standpoint. Our study supports transdiagnostic and dimensional models of neurodevelopment that transcend traditional psychiatric categories (49). Future work should focus on understanding the relation between psychopathology and brain development during early childhood (50) and incorporating measures of brain connectivity (4).

## Acknowledgments

VC was supported by a National Health and Medical Research Council (NHMRC) Investigator Grant (1177370) and a Brain and Behavior Research Foundation (NARSAD) Young Investigator Award (21660). YT was supported by the China Scholarship Council-University of Melbourne Research Scholarship. LC was supported by the Australian National Health Medical Research Council (1138711). CP was supported by a NHMRC Senior Principal Research Fellowship (ID: 1105825) and NHMRC Program Grant (ID: 1150083). AZ was supported by an NHMRC Senior Research Fellowship (ID: 1136649).

The Philadelphia Neurodevelopment Cohort sample is a publicly available data set. Support for the collection of the data sets was provided by grant RC2MH089983 awarded to Raquel Gur and RC2MH089924 awarded to Hakon Hakonarson. All subjects were recruited through the Center for Applied Genomics at The Children’s Hospital in Philadelphia. Database of Genotypes and Phenotypes study accession: phs000607.v2.p2.

