## Supplementary Material for "Brain-predicted age associates with psychopathology dimensions in youth"

**Supplementary Online Content**

### 1. Methods

#### 1.1. Participants

Participants were recruited from the Philadelphia Neurodevelopmental Cohort (PNC). Of 1598 participants for whom a structural MRI brain scan was acquired, 285 participants were excluded due to presence of severe medical conditions (*n*=74), image quality control (*n*=209) or missing clinical data (*n*=2), leaving a total of 1313 youths. This final sample was parsed into a typically developing (TD) reference group and a non-TD group for testing associations. In our final sample with MRI (i.e. after the above exclusion criteria was applied), 402 youths were identified as TD, operationalized as having no medical conditions that could affect the central nervous system (CNS), no history of psychiatric hospitalization, not currently taking psychotropic medication and not meeting criteria for psychosis spectrum symptoms (1). Ratings for medical health conditions was performed by trained personnel in the PNC team. This conservative criteria for TD resulted in a reference group that was characterised by an absence of medical (none or mild), developmental or significant psychiatric conditions for training the normative model to predict individual age. As a corollary, the remaining 911 youth were allocated to the non-TD group if they could not be confirmed to meet the above medical and psychiatric constraints, including due to missing data. This meant that the non-TD youth may, but not necessarily, have had psychosis spectrum symptoms and/or medical, including developmental (e.g. birth defects, learning difficulties) conditions that could affect the CNS. The non-TD youths were characterized by current psychotropic medication use (*n*=124), history of hospitalization (*n*=39), psychosis-spectrum symptoms (*n*=347), and medical conditions rated by the PNC study team as none/mild (*n*=550), moderate (*n*=256) and significant (*n*=50). Our proportion of TD individuals (*n*=402, 30%) is comparable to other studies using the PNC sample (27%; (2)). Supplementary analyses were undertaken to investigate the extent to which any associations between brain age and psychopathology were contingent on the operationalization of TD.

#### 1.2. Clinical assessment

As described elsewhere (3), a computerized protocol (GOASSESS) based on the Kiddie-Schedule Affective Disorder and Schizophrenia (4) was used to assess symptoms of anxiety, mood, attention deficits, eating, behavioral problems and psychosis spectrum disorders. Clinical symptoms were obtained for all individuals (n= 9498) participating in the PNC, with collateral informants used for individuals aged 8-10 years. Following Alnæs and colleagues (5), we included a total of 129 symptom items spanning 18 clinical screening categories (**Supplementary Table 2**) for each individual, excluding follow-up and conditional items. The 18 clinical screening categories include attention deficit hyperactivity disorder (ADHD), agoraphobia, conduct disorder, major depression disorder, eating disorder, generalized anxiety, mania/hypomania, obsessive-compulsive disorder, oppositional defiant disorder, panic disorder, specific phobia, post-traumatic stress disorder, separation anxiety, suicide, psychosis, psychosis prodromal symptoms as well as general probes of psychopathology. Participants with missing responses for all 129 items were excluded, resulting in a final sample comprising 9312 individuals. The remaining missing responses from participants in each item (1.8% on average) were then replaced with the nearest non-missing neighbor value based on the Euclidean distance (5).

#### 1.3. Independent component analysis

Independent component analysis (ICA) was used to estimate latent dimensions in the broad catalogue of psychopathology. We followed the same procedure as Alnæs and colleagues (5), whom previously validated this dimensional approach in the PNC. Age was regressed from all items and the residuals were submitted to ICA using the Icasso package (6) for MATLAB, which provides an implementation of the FastICA algorithm (7). Under the ICA, the matrix of residuals (symptoms $\times$ individuals), denoted by $Y$, was factorized such that $S=W\times Y$, where $W$ denotes the de-mixing matrix (components $\times$ symptoms) and $S$ denotes the estimated independent sources (components $\times$ individuals).

To estimate the de-mixing matrix, FastICA was performed with default parameter settings, which include a cubic nonlinearity and deflationary orthogonalization. Bootstrapping (n=5000) was used to sample individuals with replacement and ICA was performed independently on each bootstrapped sample with randomly chosen initial conditions. Independent components that represented a consensus across the 5000 bootstrapped samples were then delineated using agglomerative clustering with average-linkage. Clearly separated clusters indicated that the independent components were consistently and reliably estimated, despite the randomization of initial conditions and bootstrap resampling among individuals. A set of candidate models ranging from 3 to 15 components was estimated by Alnæs and colleagues (5), and the model comprising 7 components was deemed to be optimal. We therefore focused on 7 independent components in this study. In addition to the bootstrapping and randomization procedures described above, we further utilized a recently developed sampling and matching process to improve the reproducibility of the estimated independent components (8). Each component represented continuous variation along a particular dimension of psychopathology, from absence of symptoms to subthreshold psychopathology. The 7 dimensions characterized symptoms related to conduct, anxiety, obsessive-compulsive, attention, depression, bipolar and psychosis (Figure1A). Each participant was thus characterized by 7 scores that indexed the extent to which they expressed each of these putative dimensions of psychopathology. A rank-based inverse normal transformation was applied to improve the normality of the distribution of the psychopathology dimensions.

#### 1.4. T1-image acquisition and quality control

Automated quality control of T1 images was performed using two Freesurfer generated measures: the Euler number and the Mahalanobis distance of cortical thickness. The Euler number measures the topological complexity of a reconstructed cortical surface. In the context of Freesurfer, the Euler number is calculated using $2-2h$, where $h$ represents the average number of holes in the cortical reconstruction between both hemispheres (9). Optimally, Freesurfer will generate a cortical reconstruction with a Euler number of 2, meaning that it contains no holes. A study using PNC data to evaluate automated quality control of T1-weighted images found that a Euler threshold of -217 was the best predictor of poor T1 quality out of the methods tested (10). The Mahalanobis distance is a measure of the distance between a data point and the centre of the entire distribution, which can be applied to multi-dimensional data, weighting dimensions differently based on their standard deviation. The Mahalanobis distance of the cortical thickness of all brain regions was utilized as a measure of T1 quality, given that cortical thickness measurements were available for multiple regions. Values for both hemispheres were calculated separately. Visual inspection of participants with a Mahalanobis distance outside 2 standard deviations were found to have poor T1 quality and were thus excluded.

#### 1.5. T1-weighted image processing

Image processing was conducted using the Freesurfer software package (version 6.0, <http://surfer.nmr.mgh.harvard.edu/>), which consists of a volume-based and a surface-based stream (11-14). The automated volume-based stream was used to extract mean volume estimates for ICV (estimate based on the Talairach transform), lateral ventricles, and subcortical and cortical areas across the whole brain. The surface-based pipeline was used to extract cortical thickness and surface area measurements through the reconstruction of a three-dimensional cortical surface model. This includes segmentation of the pial surface and the grey/white matter boundaries for each hemisphere, using image intensity and continuity information from the MRI volume. Thickness measures were obtained by calculating the shortest distance between the grey/white matter boundary and the pial surface at vertices on a uniform triangular grid with 1mm spacing across the cortex. The surface area was obtained using the shortest distance between vertices on the white surface. Cortical parcellations and subcortical volumes were extracted for each participant and averaged across the left and right hemispheres. All regional cortical measurements (thickness, volume and area) were obtained using the Desikan-Killiany brain atlas (15).

#### 1.6. Diffusion tensor image acquisition and processing

A 3 Tesla Siemens MRI scanner was used to acquire a diffusion-weighted scan using a twice-refocused spin-echo single-shot echo-planar imaging sequence with the following parameters: TR = 8100 ms, TE = 82 ms, FOV = 240 mm2/240 mm2; Matrix = RL: 128/AP:128/Slices:70, in-plane resolution (x & y) 1.875 mm2; slice thickness = 2 mm, gap = 0; FlipAngle = 90°/180°/180°, volumes = 71, GRAPPA factor = 3, bandwidth = 2170 Hz/pixel, PE direction = AP. The sequence consisted of 64 diffusion-weighted directions with b = 1000 s/mm2 and 7 interspersed scans where b = 0 s/mm2.

Image processing was performed with FSL version 5.11, which is a part of the Functional Magnetic Resonance Imaging of the Brain (FMRIB) Software Library. Eddy currents and movement were corrected using FSL’s Eddy tool (16), which included rotation of diffusion gradients, outlier replacement (17) and slice-to-volume motion correction (18). Fractional anisotropy (FA), axial (AD), mean (MD) and radial (RD) diffusivity maps were computed by fitting a tensor model to the corrected DTI data using FSL’s DTIfit. All diffusivity maps were non-linearly normalized to the Montreal Neurological Space using FSL’s FMRIB58_FA template as the reference target. Regional estimates of each diffusion metric (FA, AD, MD, RD) were subsequently extracted by applying the John Hopkins University (JHU) parcellation atlas. This parcellation comprises 48 regional estimates of white matter tracts, resulting in a total of 192 regional white matter measures.

Prior to feature extraction, automated quality control of DTI images was performed using the temporal signal-to-noise ratio (tSNR), as validated previously in the PNC sample (19). In this study, a tSNR cut-off of 5.7 was found to be the best predictor of poor DTI quality. In our sample, 145 participants (10.4%) had a tSNR value <5.7. However, this tSNR cut-off (19) was applied to non-eddy corrected data, which is not optimal given the substantial improvements to image motion and distortion provided by Eddy. When applying the tSNR cut-off of 5.7 to eddy-corrected data, only 3 scans met this criterion. We thus applied a conservative criterion and excluded participants with the 10.4% lowest tSNR values (n=145), to be consistent with the number identified with the cut-off of 5.7 before eddy correction.

A total of 1396 participants in the PNC had acquired a diffusion-weighted scan. Of these participants, 145 were excluded due to poor tSNR (see above) and a further 58 were excluded due to major medical conditions that could affect brain function, leaving a total of 1193 participants with diffusion-weighted scans. Taking into consideration the participants that were excluded due to T1 quality control and missing clinical data (see main text and above), the sample size of individuals with *both* T1-weighted and diffusion-weighted images was 1069.

#### 1.7. Supplementary analyses on brain age gap – psychopathology associations

We evaluated the robustness of the observed associations between brain age gap and psychopathology dimensions with the following supplementary analyses;

1.7.1. The same general linear model reported in the main analyses tested whether brain age gap associated with each of the dimensions of psychopathology in the whole sample of youth (*n*=1313), rather than only the non-TD (*n*=911) youth. This analysis was performed to determine whether the observed associations remained with the inclusion of TD individuals, given that these individuals may have reported non-zero levels of psychopathology. The model was fitted independently to each of the 8 psychopathology scores.

1.7.2. We contend that our TD group provided an optimal reference for training the normative model to predict individual age, without being confounded by conditions that could affect brain developmental or health. Nevertheless, to determine whether the observed relationships between brain age gap and dimensions of psychosis, OCS and general psychopathology were influenced by the definition of our TD and non-TD groups, we examined alternative models for predicting age and the subsequent psychopathology associations. Using 10-fold cross-validation, we trained the model on the entire cohort, rather than only the TD group. An individual’s age was predicted using the model that was trained when they comprised the test fold. The 10-fold cross-validation was repeated, each time randomizing the allocation of individuals to folds, and the results were averaged across the repeats. A general linear model subsequently tested whether brain age gap associated with dimensions of psychopathology in the whole sample (see main text).

1.7.3. To determine whether certain confounds influenced the observed relationships between brain age gap and psychopathology, a general linear model was tested with inclusion of the following covariates: medical rating (0 – 3, as rated by the PNC), psychotropic medication (yes/no), IQ, and trauma-stress exposure. A cumulative measure of trauma exposure load (0-8) was computed by summing the number of endorsed traumatic-stressful events in the GOASSES (PTD001, PTD002, PTD003, PTD004, PTD006, PTD007, PTD008, PTD009), following Gur et al (20). Note that the same trauma items were included in the ICA and were thus reflected across the 7 psychopathology dimensions, particularly the “Conduct” dimension. In contrast to Gur et al (20) an index of socioeconomic status as a measure of poverty was not available in the PNC database.

A general linear model tested whether brain age gap associated with each of the dimensions of psychopathology in the non-TD youth (*n*=911). Specifically, the brain age gap of the $i$th participant was modeled as,

$$BAG_{i}=\beta_{0}+S_{i}\beta_{1}+age_{i}\beta_{2}+rating_{i}\beta_{3}+med_{i}\beta_{4}+IQ_{i}\beta_{5}+TSE_{i}\beta_{6}+ \epsilon_{i}$$

where $BAG_{i}$ and $S_{i}$ denote the residualized brain age gap and psychopathology score of the $i$th participant, respectively, $\beta$ denotes the fitted regression coefficients and $\epsilon_{i}$ is the error term. In addition, $rating_{i}$ denotes medical rating, $med_{i}$ denotes medication status, $IQ_{i}$ denotes IQ and $TSE_{i}$ denotes traumatic-stress exposure of the ith participant, respectively. The model was fitted independently to each of the 8 psychopathology scores.

#### 1.8. Associations between subcortical and cortical brain structures with psychopathology

The main analyses in this study establish an association between individual variation in psychopathology and the brain age gap. In supplementary analyses, we substituted the brain age gap with measures of cortical and subcortical morphology and tested whether they associated with psychopathology, while controlling for the effects of age and sex. In this way, we treated age as a confound, rather than explicitly modeling age as part of a normative model. Specifically, linear regression analysis was used to test whether individual variation in subcortical volumes associated with each of the dimensions of psychopathology. For each of the 7 subcortical regions, the variance in subcortical volumes explained by each of the 8 dimensions was evaluated independently, while controlling for the effect of age and sex. In particular, the subcortical volume of the $i$th participant was modeled as,

$$volume_{i}=\beta_{0}+S_{i}\beta_{1}+age_{i}\beta_{2}+sex_{i}\beta_{3}+\epsilon_{i}$$

where $S_{i}$ denotes the psychopathology score of the $i$th participant, $\beta$denotes the fitted regression coefficients and $\epsilon$ is the error term. The null hypothesis of an absence of any association between volume and psychopathology (${H_{0}: \beta}_{1}=0$) was evaluated independently for each dimension and subcortical region. All participants, TD and non-TD, were included in the analysis. Error control was achieved by enforcing a false-discovery rate (FDR) of 5% (21) across the 56 tests (7 regions × 8 psychopathology dimensions).

To test whether individual variation in surface based cortical structure (area, thickness and volume) associated with psychopathology dimensions, the above regression model was independently fitted to each vertex comprising the surface mesh. As a result, each vertex was endowed with a *t*-statistic quantifying evidence against the null hypothesis (${H_{0}: \beta}_{1}=0$). Spatially contiguous clusters of vertices were identified among the vertices with a *t*-statistic exceeding 3. Permutation testing was then used to estimate a family-wise error corrected p-value for each cluster (5000 permutations). These steps were performed using the Permutation Analysis of Linear Models (PALM) software (22). Two alternative hypotheses were tested separately; namely, a positive ($\beta_{1}>0$) and a negative ($\beta_{1}<0$) association between cortical structure and psychopathology. Error control was achieved by enforcing a FDR of 5% across the 42 tests (2 hemispheres × 7 psychopathology dimensions × 3 cortical measures).

The spin test (23) was used to test whether the areas of the cortex where the relationship between cortical structure and psychopathology was strongest/weakest were spatially consistent across the dimensions of psychopathology. The spin test was performed on the *t*-statistic maps for $\beta_{1}$ derived from the above regression model. In particular, the Pearson correlation coefficient was computed across all vertices to evaluate the strength of spatial correspondence between pairs of *t*-statistic maps. This yielded a 7 × 7 correlation matrix that quantified the spatial correspondence between all dimension pairs. To determine a p-value for each correlation coefficient, one of the *t*-statistic maps was rotated about a randomly chosen axis and a new correlation coefficient was computed. This was repeated for 1000 random rotations, thereby yielding a null distribution for the correlation coefficient. Error control was achieved by enforcing a FDR of 5% across the 21 pairs of psychopathology dimensions.

### 2. Results

#### 2.1. Sample characteristics

Demographic and clinical characteristics of the sample are presented in **Supplementary Table 1**. The mean age of the sample was 14.5 years (8-21 years) and the age distribution of males and females were equal (Kolmogorov-Smirnov: p=0.99). The estimated IQ of the sample was average (mean estimated IQ = 101.9) in relation to their age group. On average, the sample had undertaken 8.2 years of education. Demographic characteristics did not substantially vary across the psychopathology dimensions, as computed by a weighted mean (supplementary Table 1a). A breakdown of various medical and developmental conditions in the sample is shown in supplementary Table 1b.

#### 2.2. Inclusion of diffusion metrics on computation of brain age gap

After quality control, the sample size of individuals with both T1-weighted and diffusion-weighted images was 1069.

In addition to the 111 grey matter features, measures of white matter integrity were submitted as feature inputs to the brain age prediction model to determine whether the inclusion of white matter features improved brain age estimation. Measures of fractional anisotropy, mean diffusivity, radial and axial diffusivity were extracted from the left and right hemisphere using the Johns-Hopkins atlas, resulting in an additional 192 regional measures of white matter, and 303 features in total. Age was predicted from the set of 303 grey and white matter features as before (see main text). The only exception was that the samples were slightly smaller, consisting of 335 typically developing youth for the training sample, and 734 youth as the test sample, with both T1-weighted and diffusion-weighted scans that passed quality control.

Using this new set of 303 grey and white matter features, the accuracy of age prediction was reduced. Within the training set, the brain features predicted chronological age with a mean absolute error (MAE) of 1.93 years (average over 100 runs) and a correlation between chronological and predicted age of *r*=0.75, *p*<0.0001. Because the poorer accuracy of brain age estimation may have been due to the smaller training sample rather than the addition of white matter features, we additionally ran the brain age model on the reduced sample but with the original 111 grey matter measures as the feature input. In the training sample this resulted in a MAE of 1.52 years, which was comparable to the MAE of 1.49 years reported in the T1-only sample (see main text). Due to the poorer brain age estimation with the addition of white matter features, our brain age prediction model consisted of only grey matter structural indices.

#### 2.3. Supplementary analyses on brain age gap – psychopathology associations

Whole sample

In the whole sample of TD and non-TD (*n*=1313), individual variation in brain age gap was significantly associated with OCS (*t*=2.5, *p*=0.012), psychosis (*t*=3.1, *p*=0.002) and general psychopathology (*t*=4.0, *p*<0.0001). Statistics reported are the average of 100 runs. Brain age gap was not associated with the remaining psychopathology dimensions.

Training on the whole sample

When the normative model to predict age was trained on a random sample of participants (rather than the TD group), the same psychopathology dimensions significantly associated with brain age gap. Specifically, individual variation in brain age gap was significantly associated with OCS (*t*=2.4, *p*=0.02), psychosis (*t*=2.9, *p*=0.003) and general psychopathology (*t*=4.3, *p*<0.0001) (average of 100 runs). Brain age gap was not associated with the remaining psychopathology dimensions.

Impact of nuisance confounds

Medical rating, psychotropic medication, IQ and TSE variables were selected as nuisance confounds and included in the general linear model as covariates of no interest. When controlling for these covariates, brain age gap remained significantly associated with OCS (*t*=2.3, *p*=0.02), psychosis (*t*=2.9, *p*=0.004) and general psychopathology (*t*=3.97, *p*<0.0001). This indicates that the associations of brain age gap with these select psychopathology dimensions are not explained by severity of medical conditions, psychotropic medication use, IQ and exposure to traumatic events.

With respect to the covariates themselves, there was no main effect of psychotropic medication (*t*=-0.16, *p*=0.87), IQ (*t*=1.04, *p*=0.3) and TSE (*t*=0.57, *p*=0.57) on brain age gap. There was however, a significant main effect of medical rating on brain age gap (*t*=-0.16, *p*=0.02), although individuals with brain morphology that appeared *younger* than predicted by the normative model (implying a delayed pattern of development) were associated with higher medical ratings. There was also a significant interaction between TSE and age (*t*=-2.8, *p*=0.004), as demonstrated by a positive relationship between TSE and brain age gap in younger individuals, and an absence of a relationship in older individuals. When including this interaction into the GLM as a covariate, brain age gap remained significantly associated with OCS (*t*=2.03, *p*=0.04), psychosis (*t*=2.8, *p*=0.005) and general psychopathology (*t*=3.9, *p*=0.0001). TSE was significantly associated with the dimensions representing conduct (*r*=0.42, *p*<0.00001), bipolar (*r*=0.16, *p*<0.00001) and psychosis (*r*=0.28, *p*<0.00001) symptoms, as well as general psychopathology (*r*=0.45, *p*<0.00001), and was marginally associated with depression (*r*=0.06, *p*=0.05) and inattention (*r*=-0.065, *p*=0.05) dimensions. For all dimensions except inattention, greater traumatic exposure load was associated with greater symptom severity.

#### 2.4. Characterizing the relationship between dimensional constructs of psychopathology and brain structure

Linear regression analysis controlling for the effects of age and sex revealed that continuous individual variation in the dimensions characterizing anxiety, psychosis, attention and conduct were significantly related to cortical structure within circumscribed brain regions (**Supplementary** **Figure 3A**). Scoring higher on the anxiety dimension was associated with widespread reductions in cortical area and volume, and lower thickness within the temporal and occipital lobes. Higher psychosis, attention and conduct scores were associated with lower thickness in discrete lateral and medial occipital-parietal regions (psychosis), the lateral surface of the pre/post central gyrus (attention) and a discrete cluster in the frontal cortex (conduct), with individual variation in conduct additionally showing volume reductions within the cingulate, parietal and temporal regions. Peak coordinates of the clusters are presented in **Supplementary Tables 6-8**. Effect sizes were moderate (peak t-stat value 3.54 – 7.38; see supplementary tables 6-8).

Subcortical volumes were significantly associated with anxiety, conduct, psychosis and general psychopathology. Similar to the cortical surface, anxiety was associated with lower volume of all subcortical regions (all *p* <0.002), whereas conduct and psychosis were specifically associated with lower hippocampal volume (*p*=0.01 and *p*=0.009, respectively). General psychopathology was associated with lower volumes of the thalamus, caudate, hippocampus and amygdala (all *p* <0.001). Correlation coefficients, t-statistic and p-values are presented in **Supplementary Table 9.**

We next sought to determine whether the regional distribution of the aforementioned effects was unique and specific to each psychopathology dimension, or whether a common locus underpinned all, or a subset of dimensions. The spin test (see above) unveiled a significant spatial overlap in the regions where the relationship between psychopathology and cortical structure was strongest/weakest (**Supplementary** **Figure 3B & C**). Specifically, regional patterns of cortical area, thickness and volume that associated with higher scores of anxiety, conduct and psychosis were common to these three psychopathology dimensions, with spatial correlations ranging from 0.2 to 0.3. Recapitulating the overlap between these three dimensions across three distinct measures of cortical morphology attests to the robustness of the relationship (Supplementary Figure 3A). In addition, dimensions characterizing anxiety and bipolar shared a common regional pattern in area; psychosis and obsessive-compulsive, and conduct and attention, were correlated in thickness; and anxiety with attention and bipolar were correlated in volume (all *r’s*= 0.1 – 0.2). Therefore, despite the 7 psychopathology dimensions being independent (i.e. individual variation in one construct was not correlated with any another), these findings indicate that the dimensions share a partially overlapping phenotype with respect to cortical structure, particularly the dimensions characterizing anxiety, conduct and psychosis. This suggests that a single aberrant pattern in cortical structure can potentially influence multiple psychopathology constructs.

### 3. Supplementary Tables

##### **Supplementary Table 1.** Characteristics of the study sample

**a)**

|  | **Whole** | **TD** | **Conduct** | **Anxiety** | **Obsessive - Compulsive** | **Attention** | **Depression** | **Bipolar** | **Psychosis** | **General** |
| --- | --- | --- | --- | --- | --- | --- | --- | --- | --- | --- |
| Age (years) | 14.5 ± 3.4 | 14.1 ± 3.6 | 14.55 ± 3.4 | 14.47 ± 3.5 | 14.52 ± 3.3 | 14.52 ± 3.4 | 14.56 ± 3.4 | 14.53 ± 3.4 | 14.50 ± 3.3 | 14.50 ± 3.3 |
| Sex | 1.50 ± 0.5 | 1.51 ± 0.5 | 1.51 ± 0.5 | 1.50 ± 0.5 | 1.47 ± 0.5 | 1.50 ± 0.5 | 1.50 ± 0.5 | 1.49 ± 0.5 | 1.51 ± 0.5 | 1.49 ± 0.5 |
| IQ | 101.9 ± 16.4 | 104.1 ± 16.2 | 99.8 ± 16.2 | 101 ± 16.4 | 102.1 ± 16.5 | 100.9 ± 16.5 | 102.1 ± 16.7 | 102.1 ± 16.5 | 101.0 ± 16.7 | 100.1 ± 16.7 |
| Education | 8.2 ± 3.4 | 7.9 ± 3.6 | 8.2 ± 3.4 | 8.1 ± 3.3 | 8.2 ± 3.3 | 8.2 ± 3.3 | 8.2 ± 3.4 | 8.2 ± 3.4 | 8.1 ± 3.2 | 8.1 ± 3.2 |

Mean ± standard deviation reported. The whole sample consisted of n=1313 youth with both MRI and clinical (ICA) ratings and was parsed into a typically developing (TD; n=402) and non-TD (n=911) group. For the psychopathology dimensions, a weighted mean and standard deviation of each demographic variable in the non-TD group is reported. TD, typically developing; General, general (overall) psychopathology

**b)**

| **Medical conditions** | **TD (*n* = 402)** | **Non-TD (*n* = 911)** |
| --- | --- | --- |
| Abnormal or delayed childhood development | - | 202 (22.2) |
| Recurrent migraines | - | 113 (12.4) |
| Ever treated for epilepsy, convulsions, seizures | - | 25 (2.7) |
| Head injury resulting in loss of consciousness | - | 48 (5.3) |
| Birth defects (lifetime) requiring treatment | - | 75 (8.2) |
| History of lead poisoning | - | 38 (4.2) |
| Speech problem | - | 177 (19.4) |
| Vocal tics | - | 12 (1.3) |
| Motor tics | - | 33 (3.6) |
| Dyslexia | - | 100 (11) |
| Learning problem requiring treatment | - | 80 (8.8) |
| Developmental disorder | - | 17 (1.9) |
| History of pulmonary problem | 72 (17.9) | 225 (24.7) |
| History of rheumatology/joint problem | 6 (1.5) | 52 (5.7) |
| History of cardiovascular problem | 13 (3.2) | 68 (7.5) |
| History of hepatology problem | 2 (0.5) | 9 (1) |
| History of endocrinology problem | 7 (1.7) | 58 (6.4) |
| History of immunology problem | 12 (3) | 54 (5.9) |
| Psychotropic medication | - | 124 (13.6) |
| Psychiatric hospitalization | - | 39 (4.28) |
| Psychosis spectrum symptom criteria | - | 347 (38%) |

*n* (%) reported. Individual participants may have endorsed more than one condition. Psychosis spectrum symptom criteria followed that developed by Calkins et al (2014) in the PNC sample.

##### **Supplementary Table 2.** List of the 129 GOASSES items submitted to the Independent Component Analysis

| **Item** | **Variable Description** |
| --- | --- |
| **Attention Deficit Disorder** | |
| ADD011 | Did you often have trouble paying attention or keeping your mind on your school, work, chores, or other activities that you were doing? |
| ADD012 | Did you often have problems following instructions and often fail to finish school, work, or other things you meant to get done? |
| ADD013 | Did you often dislike, avoid, or put off school or homework (or any other activity requiring concentration) |
| ADD014 | Did you often lose things you needed for school or projects at home (assignments or books) or make careless mistakes in school work or other activities? |
| ADD015 | Did you often have trouble making plans, doing things that had to be done in a certain kind of order, or that had a lot of different steps? |
| ADD016 | Did you often have people tell you that you did not seem to be listening when they spoke to you or that you were daydreaming? |
| ADD020 | Did you often have difficulty sitting still for more than a few minutes at a time, even after being asked to stay seated, or did you often fidget with your hands or feet or wiggle in your seat or were you "always on the go"? |
| ADD021 | Did you often blurt out answers to other people's questions before they finished speaking or interrupt people abruptly? |
| ADD022 | Did you often join other people's conversations or have trouble waiting your turn (e.g., waiting in line, waiting for a teacher to call on you in class)? |
| **Agoraphobia** | |
| AGR001 | Have you ever been very nervous or afraid of: being in crowds (for example, a classroom, cafeteria, restaurant, or movie theater)? |
| AGR002 | Have you ever been very nervous or afraid of: going to public places (such as a store or shopping mall)? |
| AGR003 | Have you ever been very nervous or afraid of: being in an open field? |
| AGR004 | Have you ever been very nervous or afraid of: going over bridges or through tunnels? |
| AGR005 | Have you ever been very nervous or afraid of: traveling by yourself? |
| AGR006 | Have you ever been very nervous or afraid of: traveling away from home? |
| AGR007 | Have you ever been very nervous or afraid of: traveling in a car? |
| AGR008 | Have you ever been very nervous or afraid of: using public transportation like a bus or SEPTA? |
| **Conduct Disorder** | |
| CDD001 | Was there ever a time when you often did things that got you into trouble with adults like lying or stealing (something worth more than $5, from family, others, or stores)? |
| CDD002 | Did you ever skip school, stay out at night later than you were supposed to (more than 2 hours), or run away from home overnight? |
| CDD003 | Did you ever set fires, break into cars, or destroy someone else's property on purpose? |
| CDD004 | Do you have a probation officer or have you ever been on probation? |
| CDD005 | Did you often bully others (hitting, threatening or scaring someone who was younger or smaller), threaten or frighten someone on purpose, or often start physical fights with others? |
| CDD006 | Have you ever been physically cruel to an animal or person (on purpose)? |
| CDD007 | Did you ever: try to hurt someone with a weapon (a bat, brick, broken bottle, knife, or gun)? |
| CDD008 | Did you ever: threaten someone? |
| CDD009 | Conduct Disorder: Did you ever: hold someone up? |
| CDD010 | Conduct Disorder: Did you ever: attack someone to steal from them? |
| CDD011 | Did you ever: trick or threaten someone into having sex with you, or did anyone ever accuse you of making them do something sexual? |
| **Depression** | |
| DEP001 | Has there ever been a time when you felt sad or depressed most of the time? |
| DEP002 | Has there ever been a time when you cried a lot, or felt like crying? |
| DEP004 | Has there ever been a time when you felt grouchy, irritable or in a bad mood most of the time; even little things would make you mad? |
| DEP006 | Has there ever been a time when nothing was fun for you and you just weren't interested in anything? |
| **Eating Disorder** | |
| EAT001 | Was there ever a time when you felt really fat or heavy, but other people said that you were too thin? |
| EAT007 | Has there been a time when your eating was out of control - you'd eat a large amount of food in a short period of time and could not stop yourself? |
| **Generalized anxiety** | |
| GAD001 | Have you ever been a worrier? |
| GAD002 | Did you worry a lot more than most children/people your age? |
| **Mania/Hypomania** | |
| MAN001 | Have there been times when you were much more active, excited or energetic than usual, had problems sitting still, or needed to move around a lot? |
| MAN002 | Has there ever been a time when you felt so full of energy that you couldn't stop doing things and didn't get tired? |
| MAN003 | Has there ever been a time when you felt like you hardly needed sleep? |
| MAN004 | Have there been times when you kept talking a lot, couldn't stop talking, talked faster than usual, had thoughts faster than usual, or had so many ideas in your head that you could hardly keep track of them? |
| MAN005 | Have you ever had a time when you felt much more happy or excited than you usually do when there was nothing special going on? |
| MAN006 | Have you ever had a time when you felt like you could do almost anything? |
| MAN007 | Has there ever been a time when you felt unusually grouchy, cranky, or irritable; when the smallest things would make you really mad? |
| **Obsessive Compulsive Disorder** | |
| OCD001 | Have you ever been bothered by thoughts that don't make sense to you, that come over and over again and won't go away, such as concern with harming others/self? |
| OCD002 | Have you ever been bothered by thoughts that don't make sense to you, that come over and over again and won't go away, such as pictures of violent things? |
| OCD003 | Have you ever been bothered by thoughts that don't make sense to you, that come over and over again and won't go away, such as thoughts about contamination/germs/illness? |
| OCD004 | Have you ever been bothered by thoughts that don't make sense to you, that come over and over again and won't go away, such as fear that you would do something/say something bad without intending to? |
| OCD005 | Have you ever been bothered by thoughts that don't make sense to you, that come over and over again and won't go away, such as feelings that bad things that happened were your fault? |
| OCD006 | Have you ever been bothered by thoughts that don't make sense to you, that come over and over again and won't go away, such as forbidden/bad thoughts? |
| OCD007 | Have you ever been bothered by thoughts that don't make sense to you, that come over and over again and won't go away, such as need for symmetry/exactness? |
| OCD008 | Have you ever been bothered by thoughts that don't make sense to you, that come over and over again and won't go away, such as religious thoughts? |
| OCD011 | Have you ever had to do something over and over again - that would have made you feel really nervous if you couldn't do it, like: cleaning or washing (for example, your hands, house)? |
| OCD012 | Have you ever had to do something over and over again - that would have made you feel really nervous if you couldn't do it, like: counting? |
| OCD013 | Have you ever had to do something over and over again - that would have made you feel really nervous if you couldn't do it, like: checking (for example, doors, locks, ovens)? |
| OCD014 | Have you ever had to do something over and over again - that would have made you feel really nervous if you couldn't do it, like: getting dressed over and over again? |
| OCD015 | Have you ever had to do something over and over again - that would have made you feel really nervous if you couldn't do it, like: going in and out a door over and over again? |
| OCD016 | Have you ever had to do something over and over again - that would have made you feel really nervous if you couldn't do it, like: ordering or arranging things? |
| OCD017 | Have you ever had to do something over and over again - that would have made you feel really nervous if you couldn't do it, like: doing things over and over again at bedtime, like arranging the pillows, sheets, or other things? |
| OCD018 | Have you ever saved up so many things that people complained or they got in the way? |
| OCD019 | Do you feel the need to do things just right (like they have to be perfect)? |
| **Oppositional Defiance Disorder** | |
| ODD001 | Was there a time when you often did things that got you into trouble with adults such as losing your temper, arguing with or talking back to adults, or being grouchy or irritable with them? |
| ODD002 | Was there a time when you often got into trouble with adults for refusing to do what they told you to do or for breaking rules at home/school? |
| ODD003 | Did you often annoy other people on purpose or blame other people for your mistakes (excluding siblings)? |
| ODD005 | Did you ever get into trouble for getting even with other people by doing things to hurt them, telling lies about them, or messing up their things? |
| ODD006 | Were you often irritable or grouchy, or did you often get angry because you thought that things were unfair? |
| **Panic Disorder** | |
| PAN001 | Have you ever had an attack like this? |
| PAN003 | Has there ever been a time when all of a sudden you felt very, very scared or uncomfortable - and your chest hurt, you couldn't catch your breath, your heart beat very fast, you felt very shaky, and sweaty/tingly/numb in your hands or feet? |
| PAN004 | Has there ever been a time when all of a sudden, you felt that you were losing control, something terrible was going to happen, that you were going crazy, or going to die? |
| **Specific Phobia** | |
| PHB001 | Have you ever been very nervous or afraid of animals or bugs, like dogs, snakes, or spiders? |
| PHB002 | Have you ever been very nervous or afraid of being in really high places, like a roof or tall building? |
| PHB003 | Have you ever been very nervous or afraid of water or situations involving water, such as a swimming pool, lake, or ocean? |
| PHB004 | Have you ever been very nervous or afraid of storms, thunder, or lightning? |
| PHB005 | Have you ever been very nervous or afraid of doctors, needles, or blood? |
| PHB006 | Have you ever been very nervous or afraid of closed spaces, like elevators or closets? |
| PHB007 | Have you ever been very nervous or afraid of flying or airplanes? |
| PHB008 | Have you ever been very nervous or afraid of any other things or situations? |
| **Psychosis** | |
| PSY001 | Have you ever heard voices when no one was there? |
| PSY020 | Did you ever hear other sounds or noises that other people couldn't hear? |
| PSY029 | Have you ever seen visions or seen things which other people could not see? |
| PSY050 | Have you ever smelled strange odors other people could not smell? |
| PSY060 | Have you ever had strange feelings in your body like things were crawling on you or someone touching you and nothing or no one was there? |
| PSY071 | Have you ever believed in things and later found out they weren't true, like people being out to get you, or talking about you behind your back, or controlling what you do or think? |
| **Post-traumatic Stress** | |
| PTD001 | Have you ever been in a flood or a tornado or an earthquake or a hurricane or some other natural disaster where you thought you were going to die or be seriously hurt? |
| PTD002 | Have you ever been in a situation where you thought you or someone close to you was going to be killed or be hurt very badly (e.g. family violence)? |
| PTD003 | Have you ever been attacked by somebody or badly beaten? |
| PTD004 | Have you ever been very upset by someone forcing you to do something sexual? |
| PTD006 | Have you ever been threatened with a weapon? |
| PTD007 | Have you ever been in a bad accident? |
| PTD008 | Other than television or at the movies, have you ever seen or heard somebody get killed or get hurt very badly or die? |
| PTD009 | Have you ever been very upset by seeing a dead body or by seeing pictures of the dead body of somebody you knew well? |
| **General Probes** | |
| SCR001 | Have you ever talked to a counselor, psychologist, social worker, psychiatrist or some other professional about your feelings or problems with your mood or behaviors? |
| SCR006 | Are you currently taking medication because of your emotions and/or behaviors? |
| SCR007 | Have you ever had to go to a hospital and stay overnight because of problems with your mood, feelings, or how you were acting? |
| **Separation Anxiety** | |
| SEP500 | Since you were 5 years old, has there ever been a time when you had a lot of worries about your (attachment figures) and were very upset or got sick (for example, felt sick to your stomach, headaches, thrown-up) when you were away from him/her? |
| SEP508 | Has there ever been a time when you wanted to stay home from school or not go to other places (for example, sleep-overs) without your (attachment figures)? |
| SEP509 | When you knew that you were going to be away from home or (attachment figure(s)), did you get very upset and worry (e.g., when you learned (attachment figure(s)) were going on an upcoming trip or night out)? |
| SEP510 | Did you ever worry/have bad dreams about something terrible happening to you or your (attachment figures) so that you would not see them again? |
| SEP511 | Were you scared to be alone in your room (or any place in your house) or did you need your (attachment figure(s)) to stay with you while you fell asleep? |
| **Structured Interview for Prodromal Symptoms** | |
| SIP001 | TROUBLE WITH FOCUS AND ATTENTION Severity Scale |
| SIP003 | I think that I have felt that there are odd or unusual things going on that I can't explain. |
| SIP004 | I think that I might be able to predict the future. |
| SIP005 | I may have felt that there could possibly be something interrupting or controlling my thoughts, feelings, or actions. |
| SIP006 | I have had the experience of doing something differently because of my superstitions. |
| SIP007 | I think I may get confused at times whether something I experience or perceive may be real or may be just part of my imagination or dreams. |
| SIP008 | I have thought that it might be possible that other people can read my mind, or that I can read others' minds |
| SIP009 | I wonder if people may be planning to hurt me or even may be about to hurt me. |
| SIP010 | I believe that I have special natural or supernatural gifts beyond my talents and natural strengths. |
| SIP011 | I think I might feel like my mind is "playing tricks" on me. |
| SIP012 | I have had the experience of hearing faint or clear sounds of people or a person mumbling or talking when there is no one near me. |
| SIP013 | I think that I may hear my own thoughts being said out loud. |
| SIP014 | I have been concerned that I might be "going crazy." |
| SIP027 | Do people ever tell you that they can't understand you? |
| SIP028 | Do people ever seem to have difficulty understanding you? |
| SIP030 | Changes in speech, disorganized communication, tangential speech Severity Scale |
| SIP032 | Do you ever feel a loss of sense of self or feel disconnected from yourself or your life? |
| SIP033 | Has anyone pointed out to you that you are less emotional or connected to people than you used to be? |
| SIP035 | Changes in perception of self, others, or the world in general: Severity Scale |
| SIP037 | EXPRESSION OF EMOTION: Severity Scale |
| SIP038 | Within the past 6 months, are you having a harder time getting your work or schoolwork done? |
| SIP039 | Within the past 6 months, are you having a harder time getting normal activities done? |
| SIP041 | Occupational Functioning Severity Scale |
| SIP043 | Avolition Severity Scale |
| **Social Anxiety** | |
| SOC001 | Was there ever a time in your life when you felt afraid or uncomfortable or really, really shy with people, like meeting new people, going to parties, or eating or drinking, writing or doing homework in front of others? |
| SOC002 | Was there ever a time in your life when you felt afraid or uncomfortable talking on the telephone or with people your own age who you don't know very well? |
| SOC003 | Was there ever a time in your life when you felt afraid or uncomfortable when you had to do something in front of a group of people, like speaking in class? |
| SOC004 | Was there ever a time in your life when you felt afraid or uncomfortable acting, performing, giving a talk/speech, playing a sport or doing a musical performance, or taking an important test or exam (even though you studied enough)? |
| SOC005 | Was there ever a time in your life when you felt afraid or uncomfortable because you were the center of attention and were concerned something embarrassing might happen and you felt very afraid or felt uncomfortable? |
| **Suicide** | |
| SUI001 | Have you ever thought a lot about death or dying? |
| SUI002 | Have you ever thought about killing yourself? |

##### **Supplementary Table 3.** Individual GOASSES items significantly associated with brain age gap in non-TD participants

| **Item** | **Variable Description** | ***r*** | ***p*** |
| --- | --- | --- | --- |
| PSY060 | *Have you ever had strange feelings in your body like things were crawling on you or someone touching you and nothing or no one was there?* | 0.1325 | 0.0001 |
| OCD004 | *Have you ever been bothered by thoughts that don't make sense to you, that come over and over again and won't go away, such as fear that you would do something/say something bad without intending to?* | 0.1323 | 0.0001 |
| MAN007 | *Has there ever been a time when you felt unusually grouchy, cranky, or irritable; when the smallest things would make you really mad?* | 0.1299 | 0.0001 |
| MAN004 | *Have there been times when you kept talking a lot, couldn't stop talking, talked faster than usual, had thoughts faster than usual, or had so many ideas in your head that you could hardly keep track of them?* | 0.1268 | 0.0001 |
| OCD001 | *Have you ever been bothered by thoughts that don't make sense to you, that come over and over again and won't go away, such as concern with harming others/self?* | 0.1241 | 0.0002 |
| SIP013 | *I think that I may hear my own thoughts being said out loud.* | 0.1222 | 0.0002 |
| OCD002 | *Have you ever been bothered by thoughts that don't make sense to you, that come over and over again and won't go away, such as pictures of violent things?* | 0.1194 | 0.0003 |
| SIP005 | *I may have felt that there could possibly be something interrupting or controlling my thoughts, feelings, or actions.* | 0.1159 | 0.0005 |
| SEP510 | *Did you ever worry/have bad dreams about something terrible happening to you or your (attachment figures) so that you would not see them again?* | 0.1145 | 0.0005 |
| OCD005 | *Have you ever been bothered by thoughts that don't make sense to you, that come over and over again and won't go away, such as feelings that bad things that happened were your fault?* | 0.1133 | 0.0006 |
| SIP003 | *I think that I have felt that there are odd or unusual things going on that I can't explain.* | 0.1128 | 0.0006 |
| SIP006 | *I have had the experience of doing something differently because of my superstitions.* | 0.1120 | 0.0007 |
| AGR006 | *Have you ever been very nervous or afraid of: traveling away from home?* | 0.1108 | 0.0008 |
| OCD014 | *Have you ever had to do something over and over again - that would have made you feel really nervous if you couldn't do it, like: getting dressed over and over again?* | 0.1098 | 0.0009 |
| DEP004 | *Has there ever been a time when you felt grouchy, irritable or in a bad mood most of the time; even little things would make you mad?* | 0.1080 | 0.0011 |
| SIP011 | *I think I might feel like my mind is "playing tricks" on me.* | 0.1064 | 0.0013 |
| PSY020 | *Did you ever hear other sounds or noises that other people couldn't hear?* | 0.1058 | 0.0014 |
| OCD017 | *Have you ever had to do something over and over again - that would have made you feel really nervous if you couldn't do it, like: doing things over and over again at bedtime, like arranging the pillows, sheets, or other things?* | 0.1057 | 0.0014 |
| SIP007 | *I think I may get confused at times whether something I experience or perceive may be real or may be just part of my imagination or dreams.* | 0.1038 | 0.0017 |
| ADD015 | *Did you often have trouble making plans, doing things that had to be done in a certain kind of order, or that had a lot of different steps?* | 0.1030 | 0.0019 |
| EAT007 | *Has there been a time when your eating was out of control - you'd eat a large amount of food in a short period of time and could not stop yourself?* | 0.0984 | 0.0029 |
| SIP004 | *I think that I might be able to predict the future.* | 0.0962 | 0.0036 |
| SIP014 | *I have been concerned that I might be "going crazy."* | 0.0956 | 0.0039 |
| AGR001 | *Have you ever been very nervous or afraid of: being in crowds (for example, a classroom, cafeteria, restaurant, or movie theater)?* | 0.0914 | 0.0057 |
| MAN003 | *Has there ever been a time when you felt like you hardly needed sleep?* | 0.0908 | 0.0061 |
| DEP002 | *Has there ever been a time when you cried a lot, or felt like crying?* | 0.0899 | 0.0066 |
| SIP041 | *Occupational Functioning Severity Scale* | 0.0896 | 0.0068 |
| DEP006 | *Has there ever been a time when nothing was fun for you and you just weren't interested in anything?* | 0.0880 | 0.0079 |
| SOC005 | *Was there ever a time in your life when you felt afraid or uncomfortable because you were the center of attention and were concerned something embarrassing might happen and you felt very afraid or felt uncomfortable?* | 0.0878 | 0.0080 |
| SEP500 | *Since you were 5 years old, has there ever been a time when you had a lot of worries about your (attachment figures) and were very upset or got sick (for example, felt sick to your stomach, headaches, thrown-up) when you were away from him/her?* | 0.0877 | 0.0081 |
| MAN006 | *Have you ever had a time when you felt like you could do almost anything?* | 0.0875 | 0.0082 |
| MAN005 | *Have you ever had a time when you felt much more happy or excited than you usually do when there was nothing special going on?* | 0.0852 | 0.0101 |
| OCD016 | *Have you ever had to do something over and over again - that would have made you feel really nervous if you couldn't do it, like: ordering or arranging things?* | 0.0836 | 0.0116 |
| PSY029 | *Have you ever seen visions or seen things which other people could not see?* | 0.0830 | 0.0122 |
| OCD008 | *Have you ever been bothered by thoughts that don't make sense to you, that come over and over again and won't go away, such as religious thoughts?* | 0.0822 | 0.0130 |
| OCD006 | *Have you ever been bothered by thoughts that don't make sense to you, that come over and over again and won't go away, such as forbidden/bad thoughts?* | 0.0818 | 0.0135 |
| MAN001 | *Have there been times when you were much more active, excited or energetic than usual, had problems sitting still, or needed to move around a lot?* | 0.0817 | 0.0137 |

##### **Supplementary Table 4.** Weighted mean brain age gap across symptom dimensions

| **Dimensions** | **Brain age gap weighted mean (years)** | **Brain age gap weighted mean (days)** | **Brain age gap SEM** | **UCI** | **LCI** |
| --- | --- | --- | --- | --- | --- |
| **Conduct** | 0.0694 | 25.32 | 0.067 | 0.201 | -0.062 |
| **Anxiety** | 0.0549 | 20.04 | 0.0664 | 0.185 | -0.075 |
| **Obsessive-Compulsive** | 0.1819 | 66.39 | 0.0689 | **0.317** | **0.047** |
| **Attention** | 0.0232 | 8.46 | 0.0672 | 0.155 | -0.109 |
| **Depression** | 0.0642 | 23.44 | 0.0675 | 0.197 | -0.068 |
| **Bipolar** | 0.0461 | 16.84 | 0.0668 | 0.177 | -0.085 |
| **Psychosis** | 0.1417 | 51.71 | 0.0692 | **0.277** | **0.006** |
| **General** | 0.2073 | 75.68 | 0.0686 | **0.342** | **0.073** |

##### **Supplementary Table 5.** List of clinical items in the GOASSESS used for OCD and Psychosis lifetime groupings

| **Variable Name** | **Item question** |
| --- | --- |
| **Obsessive Compulsive Disorder** | |
| OCD001 | Have you ever been bothered by thoughts that don't make sense to you, that come over and over again and won't go away, such as concern with harming others/self? |
| OCD002 | Have you ever been bothered by thoughts that don't make sense to you, that come over and over again and won't go away, such as pictures of violent things? |
| OCD003 | Have you ever been bothered by thoughts that don't make sense to you, that come over and over again and won't go away, such as thoughts about contamination/germs/illness? |
| OCD004 | Have you ever been bothered by thoughts that don't make sense to you, that come over and over again and won't go away, such as fear that you would do something/say something bad without intending to? |
| OCD005 | Have you ever been bothered by thoughts that don't make sense to you, that come over and over again and won't go away, such as feelings that bad things that happened were your fault? |
| OCD006 | Have you ever been bothered by thoughts that don't make sense to you, that come over and over again and won't go away, such as forbidden/bad thoughts? |
| OCD007 | Have you ever been bothered by thoughts that don't make sense to you, that come over and over again and won't go away, such as need for symmetry/exactness? |
| OCD008 | Have you ever been bothered by thoughts that don't make sense to you, that come over and over again and won't go away, such as religious thoughts? |
| OCD011 | Have you ever had to do something over and over again - that would have made you feel really nervous if you couldn't do it, like: cleaning or washing (for example, your hands, house)? |
| OCD012 | Have you ever had to do something over and over again - that would have made you feel really nervous if you couldn't do it, like: counting? |
| OCD013 | Have you ever had to do something over and over again - that would have made you feel really nervous if you couldn't do it, like: checking (for example, doors, locks, ovens)? |
| OCD014 | Have you ever had to do something over and over again - that would have made you feel really nervous if you couldn't do it, like: getting dressed over and over again? |
| OCD015 | Have you ever had to do something over and over again - that would have made you feel really nervous if you couldn't do it, like: going in and out a door over and over again? |
| OCD016 | Have you ever had to do something over and over again - that would have made you feel really nervous if you couldn't do it, like: ordering or arranging things? |
| OCD017 | Have you ever had to do something over and over again - that would have made you feel really nervous if you couldn't do it, like: doing things over and over again at bedtime, like arranging the pillows, sheets, or other things? |
| OCD018 | Have you ever saved up so many things that people complained or they got in the way? |
| OCD019 | Do you feel the need to do things just right (like they have to be perfect)? |
| **Psychosis** | |
| PSY001 | Have you ever heard voices when no one was there? |
| PSY020 | Did you ever hear other sounds or noises that other people couldn't hear? |
| PSY029 | Have you ever seen visions or seen things which other people could not see? |
| PSY050 | Have you ever smelled strange odors other people could not smell? |
| PSY060 | Have you ever had strange feelings in your body like things were crawling on you or someone touching you and nothing or no one was there? |
| PSY071 | Have you ever believed in things and later found out they weren't true, like people being out to get you, or talking about you behind your back, or controlling what you do or think? |

##### **Supplementary Table 6.** Cortical area clusters significantly associated with psychopathology

| **Dimension** | **Hemisphere** | **Cluster Index** | **Peak Tstat** | **Peak coordinates** | | | **Size (mm^2^)** | **NVtxs** | **Annotation** |
| --- | --- | --- | --- | --- | --- | --- | --- | --- | --- |
|  |  |  |  | Tal X | Tal Y | Tal Z |  |  |  |
| **Conduct** | Left | C1 | 4.53 | 15.7 | -27.8 | 71.9 | 438.39 | 1533 | postcentral |
| **Anxiety** | Left | C1 | 7.04 | -21.4 | -18.7 | -62.2 | 36080.20 | 94099 | inferiortemporal |
|  | Right | C1 | 6.52 | 24.7 | -26.8 | -58.0 | 38753.86 | 100419 | inferiortemporal |

##### **Supplementary Table 7.** Cortical thickness clusters significantly associated with psychopathology

| **Dimension** | **Hemisphere** | **Cluster Index** | **Peak Tstat** | **Peak coordinates** | | | **Size (mm^2^)** | **NVtxs** | **Annotation** |
| --- | --- | --- | --- | --- | --- | --- | --- | --- | --- |
|  |  |  |  | Tal X | Tal Y | Tal Z |  |  |  |
| **Conduct** | Left | C1 | 4.41 | -2.4 | -42.4 | -49.2 | 284.97 | 763 | fusiform |
|  |  | C2 | 4.96 | -26.7 | 12.6 | 34.0 | 90.24 | 313 | precentral |
| **Anxiety** | Left | C1 | 5.92 | 23.6 | -56.5 | -14.7 | 130.89 | 439 | precuneus |
|  |  | C2 | 4.46 | -8.5 | -30.2 | -55.3 | 74.92 | 166 | fusiform |
|  | Right | C1 | 4.63 | -4.7 | -86.0 | 17.8 | 112.85 | 258 | superiorparietal |
|  |  | C2 | 5.10 | 39.5 | 2.6 | -41.6 | 85.76 | 252 | superiortemporal |
|  |  | C3 | 3.81 | -22.5 | -43.9 | -19.2 | 83.83 | 299 | isthmuscingulate |
|  |  | C4 | 4.56 | 8.6 | -37.7 | -53.0 | 80.23 | 154 | fusiform |
|  |  | C5 | 4.61 | -13.5 | -44.0 | -33.4 | 77.87 | 236 | lingual |
|  |  | C6 | 4.97 | 26.7 | -68.1 | -22.5 | 65.08 | 134 | lateraloccipital |
| **Attention** | Left | C1 | 4.14 | -16.3 | -16.8 | 49.9 | 99.48 | 343 | postcentral |
|  |  | C2 | 4.33 | -28.9 | 12.1 | 31.4 | 66.36 | 236 | precentral |
| **Psychosis** | Left | C1 | 4.79 | 25.1 | -89.0 | 13.4 | 311.57 | 852 | cuneus |
|  |  | C2 | 4.68 | 26.7 | -60.5 | 3.5 | 220.51 | 647 | precuneus |
|  |  | C3 | 4.90 | -18.4 | 0.1 | 47.5 | 147.18 | 698 | precentral |
|  |  | C4 | 5.49 | -35.9 | -8.8 | 28.1 | 137.59 | 501 | postcentral |
|  |  | C5 | 4.65 | 12.9 | -40.6 | -35.2 | 104.59 | 297 | parahippocampal |
|  |  | C6 | 3.9 | -12.0 | -17.0 | 54.9 | 92.4 | 336 | postcentral |
|  |  | C7 | 4.75 | 25.9 | -35.9 | 65.9 | 73.76 | 248 | precuneus |
|  |  | C8 | 3.70 | 27.8 | -20.5 | 67.5 | 70.03 | 231 | paracentral |
|  | Right | C1 | 4.24 | -25.0 | -71.8 | 0.3 | 277.78 | 727 | cuneus |
|  |  | C2 | 4.32 | 24.7 | -71.9 | -23.0 | 190.02 | 384 | lateraloccipital |
|  |  | C3 | 4.95 | -29.5 | -21.7 | 69.4 | 150.58 | 555 | paracentral |
|  |  | C4 | 4.80 | -14.4 | -61.6 | -35.1 | 143.01 | 305 | lingual |
|  |  | C5 | 4.16 | 21.4 | -42.5 | -52.5 | 122.11 | 229 | fusiform |
|  |  | C6 | 4.42 | -8.1 | -87.4 | 18.3 | 94.89 | 224 | superiorparietal |
|  |  | C7 | 4.09 | -22.8 | -78.2 | -23.7 | 80.82 | 200 | lingual |
|  |  | C8 | 3.88 | 1.4 | -69.7 | 33.7 | 63.63 | 176 | inferiorparietal |

##### **Supplementary Table 8.** Cortical volume clusters significantly associated with psychopathology

| **Dimension** | **Hemisphere** | **Cluster Index** | **Peak Tstat** | **Peak coordinates** | | | **Size (mm^2^)** | **NVtxs** | **Annotation** |
| --- | --- | --- | --- | --- | --- | --- | --- | --- | --- |
|  |  |  |  | Tal X | Tal Y | Tal Z |  |  |  |
| **Conduct** | Left | C1 | 4.62 | 36.0 | 34.6 | 19.5 | 163.80 | 274 | caudal anterior cingulate |
|  |  | C2 | 4.16 | 36.5 | 68.8 | -10.7 | 152.69 | 207 | rostral anterior cingulate |
|  |  | C3 | 4.62 | 10.5 | -103.0 | -3.7 | 122.44 | 277 | lateral occipital |
|  |  | C4 | 4.80 | 14.9 | -28.7 | 71.6 | 102.61 | 363 | postcentral |
|  |  | C5 | 4.96 | 23.4 | -33.1 | 70.0 | 95.64 | 349 | paracentral |
|  |  | C6 | 4.02 | -37.4 | -14.5 | 10.1 | 77.47 | 269 | supramarginal |
|  |  | C7 | 3.77 | -25.1 | -10.6 | -4.2 | 73.15 | 413 | supramarginal |
|  |  | C8 | 4.38 | 24.7 | -52.2 | -13.9 | 67.44 | 274 | isthmus cingulate |
|  | Right | C1 | 4.71 | -34.2 | 11.5 | 27.7 | 217.52 | 418 | posterior cingulate |
|  |  | C2 | 4.10 | -26.0 | -84.5 | 7.7 | 96.12 | 237 | cuneus |
|  |  | C3 | 3.73 | 37.0 | 10.7 | -56.6 | 84.87 | 197 | middle temporal |
|  |  | C4 | 4.11 | 4.5 | -3.4 | 58.1 | 81.33 | 303 | precentral |
|  |  | C5 | 3.69 | -7.5 | -45.1 | 61.9 | 78.92 | 249 | superior parietal |
|  |  | C6 | 3.71 | 28.2 | -56.8 | -1.0 | 71.49 | 203 | inferior parietal |
| **Anxiety** | Left | C13 | 4.58 | 23.0 | -62.4 | 54.5 | 202.39 | 763 | superior parietal |
|  |  | C14 | 4.62 | -8.0 | 72.9 | -38.7 | 199.94 | 594 | parsorbitalis |
|  |  | C15 | 4.68 | 9.4 | 58.2 | -51.2 | 194.16 | 422 | lateral orbitofrontal |
|  |  | C16 | 5.65 | -36.7 | 2.0 | -58.9 | 167.63 | 370 | middle temporal |
|  |  | C17 | 3.99 | 18.7 | 83.1 | -48.2 | 163.53 | 313 | lateral orbitofrontal |
|  |  | C18 | 6.17 | -0.2 | -5.1 | -58.0 | 153.18 | 588 | parahippocampal |
|  |  | C19 | 3.75 | -26.1 | -69.3 | -33.3 | 118.33 | 253 | inferiortemporal |
|  |  | C20 | 4.67 | 28.4 | 72.3 | 36.1 | 107.59 | 210 | superiorfrontal |
|  |  | C21 | 4.44 | 28.7 | -17.6 | 64.8 | 105.59 | 299 | paracentral |
|  |  | C22 | 3.94 | 29.8 | 6.3 | 53.4 | 101.86 | 232 | paracentral |
|  |  | C23 | 4.24 | 18.7 | -56.3 | -29.1 | 98.31 | 277 | lingual |
|  |  | C24 | 4.67 | -31.0 | 0.1 | -22.9 | 95.24 | 293 | transversetemporal |
|  |  | C25 | 3.61 | 25.5 | -88.9 | 12.3 | 90.58 | 252 | cuneus |
|  |  | C26 | 4.12 | 4.9 | -49.8 | 58.1 | 90.53 | 268 | superior parietal |
|  |  | C27 | 3.54 | -7.0 | 58.7 | 34.3 | 88.48 | 194 | caudal middle frontal |
|  |  | C28 | 4.88 | -22.8 | 23.6 | -68.6 | 82.24 | 223 | middle temporal |
|  | Right | C1 | 6.49 | 8.9 | -21.3 | -58.6 | 10206.94 | 23957 | fusiform |
|  |  | C2 | 7.08 | 18.2 | 15.2 | -8.6 | 3352.54 | 12225 | postcentral |
|  |  | C3 | 4.92 | -12.0 | 63.7 | -51.8 | 612.45 | 1435 | lateral orbitofrontal |
|  |  | C4 | 5.10 | -3.6 | -13.5 | 64.6 | 602.15 | 2082 | postcentral |
|  |  | C5 | 4.25 | -33.7 | 80.3 | -23.9 | 337.24 | 452 | rostral anterior cingulate |
|  |  | C6 | 4.20 | -25.6 | 100.4 | 1.9 | 248.68 | 546 | superiorfrontal |
|  |  | C7 | 4.40 | -17.5 | -40.4 | 69.4 | 248.58 | 866 | superior parietal |
|  |  | C8 | 4.98 | -12.3 | 68.6 | 41.7 | 245.45 | 526 | superior frontal |
|  |  | C9 | 4.53 | -3.8 | -46.5 | 57.4 | 204.70 | 618 | superior parietal |
|  |  | C10 | 4.72 | 7.2 | 62.5 | 31.4 | 179.44 | 450 | caudal middle frontal |
|  |  | C11 | 4.24 | 16.9 | 45.9 | 30.3 | 167.80 | 428 | caudal middle frontal |
|  |  | C12 | 4.12 | -30.8 | -18.3 | 62.1 | 157.14 | 475 | paracentral |
|  |  | C13 | 4.29 | 35.4 | -48.8 | -33.5 | 156.37 | 448 | middletemporal |
|  |  | C14 | 5.35 | 32.2 | 26.9 | 16.5 | 152.20 | 465 | precentral |
|  |  | C15 | 4.45 | -4.3 | 102.7 | -8.8 | 140.98 | 264 | rostral middle frontal |
|  |  | C16 | 4.94 | -28.9 | 88.3 | 15.6 | 129.78 | 270 | superiorfrontal |
|  |  | C17 | 4.88 | -17.6 | -106.8 | -14.6 | 119.57 | 358 | lateraloccipital |
|  |  | C18 | 4.74 | -5.1 | 101.0 | -34.2 | 117.06 | 229 | rostral middle frontal |
|  |  | C19 | 4.67 | -28.5 | 92.5 | -37.1 | 102.69 | 199 | medial orbito frontal |
|  |  | C20 | 4.41 | -29.0 | 35.9 | 56.4 | 93.91 | 223 | superiorfrontal |
|  |  | C21 | 5.12 | 0.9 | -4.0 | -57.7 | 91.65 | 311 | parahippocampal |
|  |  | C22 | 4.23 | -30.6 | 16.5 | 53.4 | 89.77 | 176 | superiorfrontal |
|  |  | C23 | 3.87 | -18.5 | 49.3 | 55.9 | 84.07 | 194 | superiorfrontal |
|  |  | C24 | 4.889 | 24.3 | 30.6 | -56.0 | 82.26 | 243 | superiortemporal |
|  |  | C25 | 4.292 | 21.6 | -6.9 | 43.2 | 77.4 | 303 | postcentral |
|  |  | C26 | 4.279 | 38.4 | -31.1 | -0.5 | 76.73 | 289 | bankssts |
|  |  | C27 | 3.668 | -30.0 | -53.6 | 49.5 | 69.35 | 237 | precuneus |
| **Psychosis** | Right | C1 | 4.26 | 6.1 | -97.3 | -24.6 | 173.53 | 345 | lateraloccipotal |
|  |  | C2 | 4.72 | -21.5 | 104.9 | -28.3 | 98.44 | 214 | frontalpole |
|  |  | C3 | 4.90 | 24.6 | 39.9 | 18.3 | 91.08 | 237 | precentral |

##### **Supplementary Table 9.** Subcortical volumes significantly associated with psychopathology

|  | **Thalamus** | | | **Caudate** | | | **Putamen** | | | **Pallidum** | | | **Hippocampus** | | | **Amygdala** | | | **Nucleus Acc** | | |
| --- | --- | --- | --- | --- | --- | --- | --- | --- | --- | --- | --- | --- | --- | --- | --- | --- | --- | --- | --- | --- | --- |
|  | *r* | *t* | *p* | *r* | *t* | *p* | *r* | *t* | *p* | *r* | *t* | *p* | *r* | *t* | *p* | *r* | *t* | *p* | *r* | *t* | *p* |
| **Conduct** |  |  |  |  |  |  |  |  |  |  |  |  | -0.07 | -2.54 | 0.01 |  |  |  |  |  |  |
| **Anxiety** | -0.14 | -5.12 | <.001 | -0.09 | -3.30 | 0.001 | -0.14 | -5.03 | <.001 | -0.08 | -3.05 | 0.002 | -0.14 | -5.12 | <.001 | -0.12 | -4.52 | <.001 | -0.09 | -3.17 | 0.002 |
| **Psychosis** |  |  |  |  |  |  |  |  |  |  |  |  | -0.07 | -2.61 | 0.009 |  |  |  |  |  |  |
| **General** | -0.12 | -4.26 | <.001 | -0.11 | -4.07 | <.001 |  |  |  |  |  |  | -0.10 | -3.62 | <.001 | **-**0.10 | -3.64 | <.001 |  |  |  |

### 4. Supplementary Figures

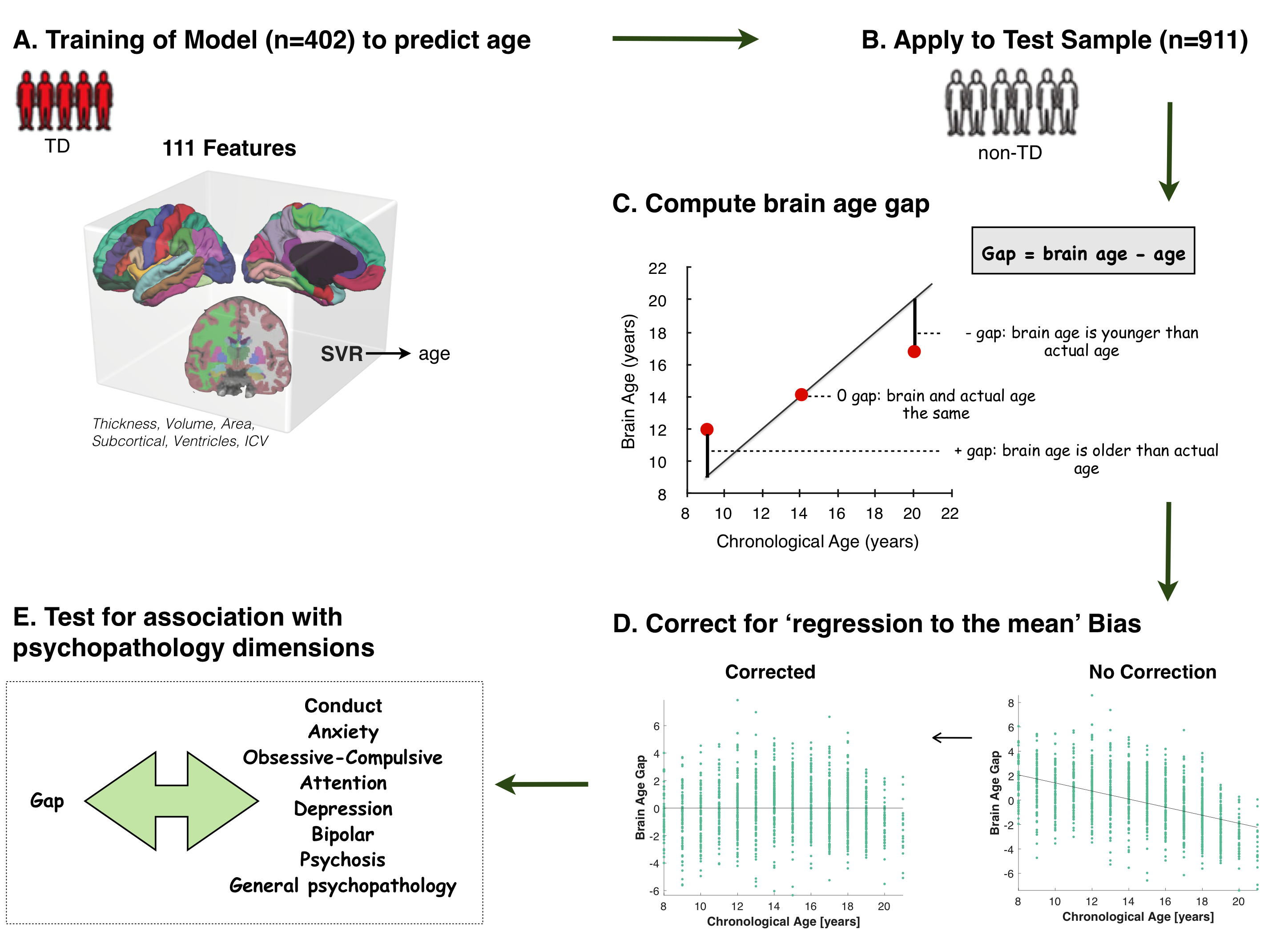

##### **Supplementary Figure 1**. Method for predicting brain age and association with psychopathology.

(**A**). A linear support vector regression (SVR) with 10-fold cross-validation was trained on 402 TD individuals to predict individual age based on a feature space comprising 111 gray matter features. Morphological features consisted of regional estimates of cortical surface area, thickness and volume based on the Desikan-Killiany parcellation atlas, 7 subcortical volumes, lateral ventricle volume and intra-cranial volume. (**B**). The SVR model trained on all TD youths was subsequently applied to predict age of each non-TD individual. (**C**). Brain age gap was calculated for each individual by subtracting chronological age from the brain-predicted age. (**D**). Chronological age was correlated with brain age gap (*r*=0.495, *p*<0.0001), and was thus regressed from brain age gap and the resulting residuals were used in all subsequent analyses (24). (**E**). A general linear model was performed to test whether individual variation in brain age gap explained variation in 7 subclinical symptoms related to conduct, anxiety, obsessive-compulsive, attention, depression, bipolar and psychosis, as well as a measure of general psychopathology.

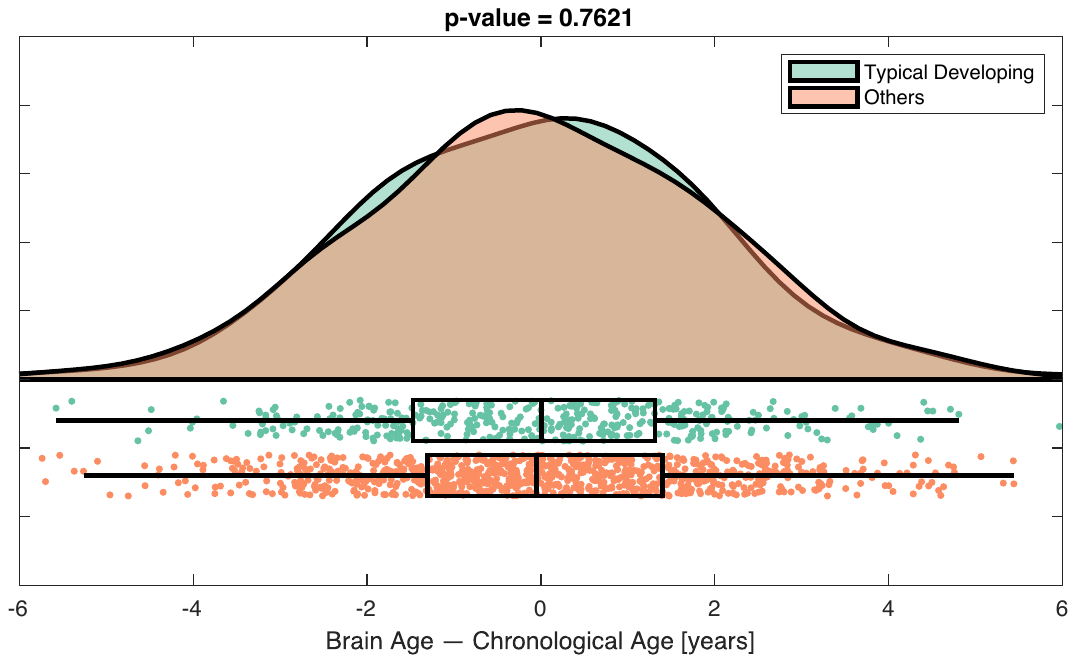

##### **Supplementary Figure 2**. Comparison of brain age gap between typically developing and non-typically developing youth.

There was no significant difference in brain age gap between the training sample (typically developing youth; n=402) and the test sample (non-typically developing youth; n=911)

.

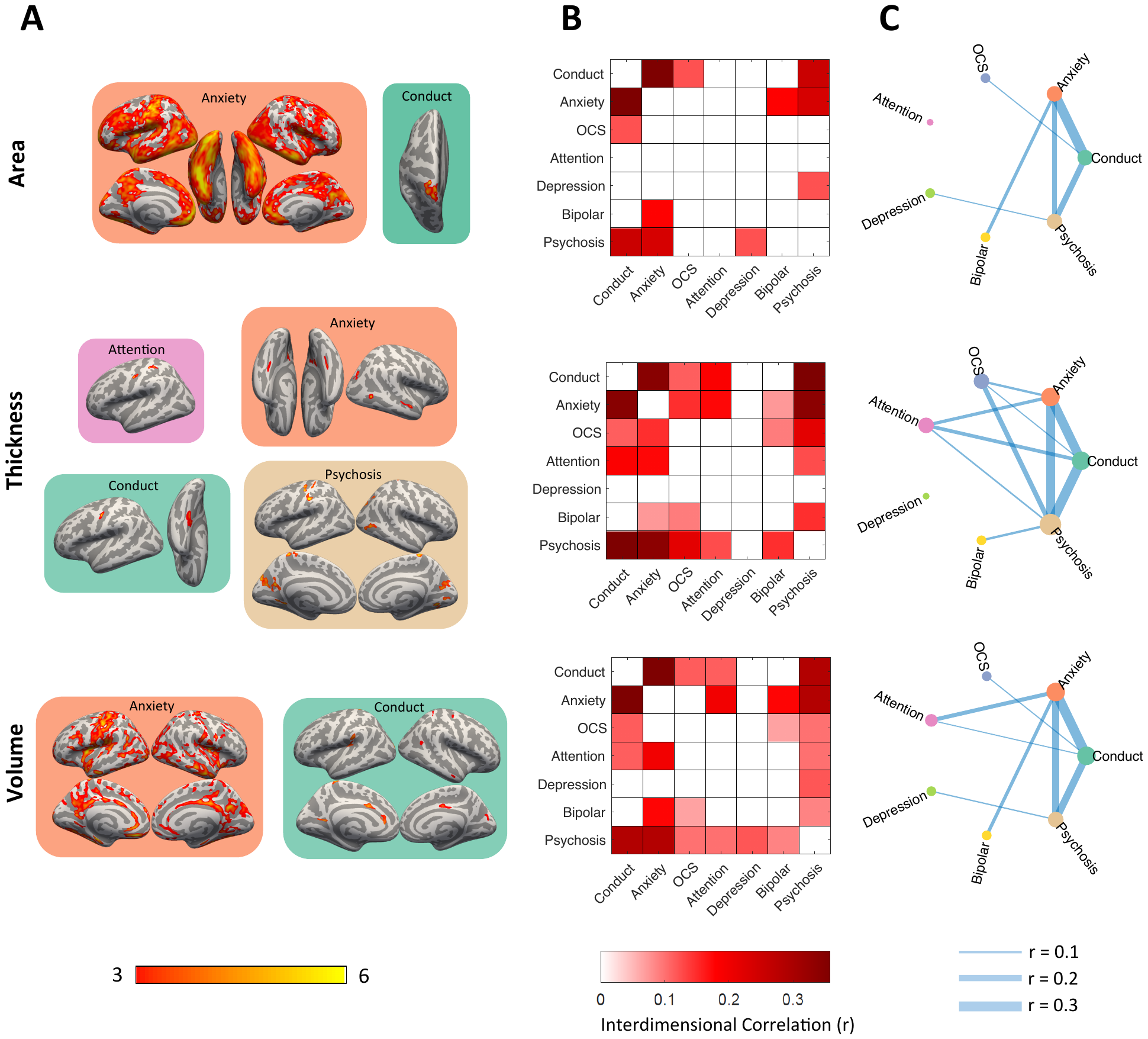

##### **Supplementary Figure 3**. Relationship between dimensions of psychopathology and cortical grey matter structure

(**A**). Surface-based analyses controlling for the effects of age and sex revealed that the dimensions characterizing anxiety, psychosis, attention and conduct were significantly related to lower cortical structure within circumscribed regions. (**B, C**). A spin test revealed a significant spatial overlap in the regions where the relationship between psychopathology and cortical structure was strongest/weakest. In particular, the dimensions characterizing anxiety, conduct and psychosis shared a common regional pattern that was seen across all three structural metrics - area, thickness and volume.
